# Neuronal TIMP2 regulates hippocampus-dependent plasticity and extracellular matrix complexity

**DOI:** 10.1101/2022.12.28.522138

**Authors:** Ana Catarina Ferreira, Brittany M. Hemmer, Sarah M. Philippi, Hanxiao Liu, Jeffrey D. Zhu, Tatyana Kareva, Tim Ahfeldt, Merina Varghese, Patrick R. Hof, Joseph M. Castellano

## Abstract

The functional output of the hippocampus, a brain region subserving memory processes, depends on highly orchestrated cellular and molecular processes that regulate synaptic plasticity throughout life. The structural requirements of such plasticity and molecular processes involved in this regulation are poorly understood. Specific molecules, including tissue inhibitor of metalloproteinases-2 (TIMP2) have been implicated in processes of plasticity in the hippocampus, a role that decreases with brain aging as expression is lost. Here, we report that TIMP2 is highly expressed by neurons within the hippocampus and its loss drives changes in cellular programs related to adult neurogenesis and dendritic spine turnover with corresponding impairments in hippocampus-dependent memory. Consistent with the accumulation of ECM in the hippocampus we observe with aging, we find that TIMP2 acts to reduce accumulation of extracellular matrix (ECM) around synapses in the hippocampus. Moreover, its removal results in hindrance of newborn neuron migration through a denser ECM network. A novel conditional TIMP2 KO mouse reveals that neuronal TIMP2 regulates adult neurogenesis, accumulation of ECM, and ultimately hippocampus-dependent memory. Our results define a mechanism whereby hippocampus-dependent function is regulated by TIMP2 and its interactions with the ECM to regulate diverse processes associated with synaptic plasticity.

## Introduction

A variety of cellular, molecular, and functional changes have now been documented in the brain as aging progresses, leaving it susceptible to diseases like Alzheimer’s disease. Given that aging serves as the major risk factor for many neurological disorders, aging processes have increasingly been considered to be valid therapeutic targets, especially as the aged population continues to increase. Growing evidence in animal models indicates that the aged brain can be rejuvenated through manipulation of the systemic environment, including through young plasma transfer or sharing of young blood through heterochronic parabiosis^1–4^. Aged mice exposed to young blood exhibit improved synaptic plasticity and cognitive performance^2^. We and others have begun to identify factors present in young blood that confer brain rejuvenation phenotypes in aged mice^3, 4^. Among those, we showed that tissue inhibitor of metalloproteinases-2 (TIMP2), a protein elevated early in life, revitalizes function of the aged mouse hippocampus^4^. Despite demonstrating that this factor is critical for young plasma-mediated memory improvements in aged mice^4^, little is known of the cellular and molecular details of how TIMP2 regulates function within the hippocampus.

TIMP2 is a secreted protein of approximately 21-kDa in size primarily known for its role in regulating peripheral extracellular matrix (ECM) remodeling^5^ through its unique dual roles in regulating the activation of latent matrix metalloproteinases (MMPs), including MMP2, to proteolytically active forms, and the inhibition of MMP activity^6, 7^. Through these roles, TIMP2 has been shown to play a key role in the maintenance of ECM network structure in many tissues with myriad disease implications, including heart failure^8^, cancer^9^, and ischemic stroke^10^, among others. While early reports described expression of TIMP2 from development into adulthood^11^, and its role in regulating cortical neuronal differentiation^12^ and subtle motor phenotypes^13^, few studies have examined the biological role of TIMP2 in the adult brain and how it modulates function of the hippocampus. While we recently demonstrated that TIMP2 is critical in revitalizing overall hippocampus-dependent cognitive function in aged mice^4^, the biological role of TIMP2 expressed within the hippocampus has not been studied. TIMP2 is part of a small family of tissue metalloproteinase inhibitors that include TIMP1, TIMP3, and TIMP4, all of which perform broad MMP targeting functions but in diverse contexts in the brain, including in myelin repair^14^, neurovascular regulation^15^, or in regulation of the tumor microenvironment^16^. Interestingly, TIMP2 shares only ∼40-50% sequence homology with other TIMP family members and is particularly expressed in the brain and enriched specifically in hippocampus relative to other TIMPs. Since levels of TIMP2 decline in the hippocampus with age^4^, understanding the function of this local source will be important to understand its normal function and position it for putative therapies in age-related brain disorders. Moreover, we sought to understand the extent to which ECM regulation is related to disparate processes of plasticity to regulate complex behavioral outputs.

Here we define a mechanism by which TIMP2 regulates ECM components in the hippocampus in a manner that regulates processes linked to plasticity and memory. We find that TIMP2 is predominantly expressed by neurons in the hippocampus and is present at high levels in the brain extracellular space adjacent to these cells. Deletion of TIMP2 induces transcriptomic changes in the hippocampus that are consistent with processes related to synaptic plasticity and adult neurogenesis. We find that loss of TIMP2 results in fewer dendritic spines in dentate gyrus (DG) granule cells, decreased adult DG neurogenesis along several stages, and impaired memory in various hippocampus-dependent tasks. Loss of TIMP2 also causes an accumulation of ECM proteins, particularly around synapses within the DG, and impaired migration of neuroblasts within the neurogenic niche, reflecting dysregulated ECM turnover and mimicking ECM accumulation seen in the aged brain. Finally, we developed a mouse model for conditional deletion of TIMP2 from neurons that exhibits the functional phenotypes observed in TIMP2 KO mice, further arguing for a role of the neuronal pool of TIMP2 in shaping plasticity-related function of the hippocampus. Our findings indicate molecular mechanisms through which neuronal TIMP2 remodeling of the ECM promotes synapse plasticity and is required for memory, with functional implications for neurogenic rejuvenation.

## Materials and Methods

See Supplementary Information for additional details.

### Animals

All animal procedures were performed in accordance with the National Institutes of Health Guide for Care and Use of Laboratory Animals and the Icahn School of Medicine at Mount Sinai Institutional Animal Care and Use Committee. TIMP2 knockout (KO) (Jackson Laboratory) and WT female mice or Syn^Cre/+^; TIMP2^fl/fl^ and TIMP2^fl/fl^ mice (generated as described in Supplementary Information) were used at 2-3 months or 5-6 months of age unless indicated otherwise. Littermate controls were used for all experiments. Mice were maintained on a 12-hour light/dark cycle at constant temperature (23°C) with *ad libitum* access to food and water.

### Bulk RNA-sequencing

RNA-sequencing was performed on hippocampi dissected from WT and TIMP2 KO male and female mice. Quality of extracted RNA (RNeasy Mini Kit, Qiagen) was measured by Agilent TapeStation Bioanalyzer (Agilent Technologies), and all samples exhibited RNA Integrity Number (RIN) > 8. cDNA libraries were prepared with poly(A) selection and sequenced using Illumina Hiseq (2x150bp paired-end) (Genewiz). At least 25M clean reads were generated from each sample and mapped to the *Mus musculus* GRCm38 reference genome available on ENSEMBL, using STAR aligner (v.2.5.2b). After extraction of gene hit counts, the hit counts table was used for downstream differential expression analysis using DESeq2 software. The Wald test was used to generate *P*-values and log2 fold changes. Differentially expressed genes (DEGs; *P*<0.05) were used for Gene Set Enrichment Analysis (GSEA; https://www.gsea-msigdb.org/gsea/index.jsp). Volcano plot was generated with R v4.1.2.

### Animal procedures

Mice were injected intraperitoneally (i.p.) once with 150 mg/kg of 5-bromo-2′-deoxyuridine (BrdU; Sigma) 24 hours before sacrifice for cell proliferation assessment, or daily for 5 days, followed by sacrifice/perfusion 28 days after initial injection for cell fate analysis. *In vivo* microdialysis proceeded according to previous methods^17^ with some adaptations (described in Supplementary Information).

### Immunohistochemistry

For all immunohistochemistry experiments, mice were anesthetized with a cocktail of ketamine (90 mg/kg) and xylazine (10 mg/kg) and transcardially perfused with ice-cold 0.9% saline, and brains were postfixed in 4% paraformaldehyde and preserved with 30% sucrose before sectioning at 40-µm on a freezing-sliding microtome (SM2010R, Leica). Sections were incubated overnight with indicated primary antibodies, followed by fluorescent secondary antibodies. Image processing was performed with LSM 780 confocal microscope (Zeiss) using 40x/1.4 Oil DIC objective. Four equally-spaced sections per mouse were used to count the total number of positive cells within DG (subgranular zone and hilar subregions) using stereological principles. All counts were performed using FIJI in a blinded fashion, according to similar methods^4^.

### Aggrecan and Homer1 puncta quantification

Quantification of the number of puncta by super-resolution microscopy proceeded according to similar methods^18^. Images were acquired by confocal imaging using LSM 880 with AiryScan in super-resolution mode (Zeiss) set with a 63X/1.4 Oil DIC objective with 5X optical zoom. Aggrecan puncta were quantified with Puncta Analyzer plugin^19^ in ImageJ, and thresholding was applied equally across images. Colocalization of co-stained Aggrecan and Homer1 puncta was analyzed with the Puncta Analyzer plugin with a minimum pixel specification of 4. Three images in the molecular layer of the DG were averaged per mouse for analysis.

### Dendritic spine analysis by iontophoretic dye injections

Tissue was processed according to similar work^20^, and then coronal sections were incubated in 250 ng/ml DAPI to enable DG identification. Sections were mounted on nitrocellulose membrane filters, immersed in ice-cold PB, and DG granule cells were iontophoretically injected with 5% Lucifer Yellow (Invitrogen) under a direct current of 3-8 nA until the dye filled distal ends of the dendrites, and then sections were mounted between spacers placed on gelatin-coated glass slides (#22-214-320, Thermo Fisher). Images of dendritic segments from DG granule cells at the suprapyramidal blade were acquired on a Zeiss LSM 780 confocal microscope using a 100x/1.46 Oil DIC M27 Plan-Apochromat objective, and stacks were acquired at 512 x 512-pixel resolution with a Z-step of 0.1 µm, optical zoom of 3.3x, pinhole setting of 1 Airy Unit, and optimal settings for gain and offset. Three z-stacks were imaged from each neuron. Confocal stacks were deconvolved using an iterative blind deconvolution algorithm (AutoQuant X, vX3.0.1, MediaCybernetics). Deconvolved stacks were analyzed using Neurolucida 360 (v2019.2.1; MBF Bioscience) for semi-automated reconstruction to determine spine density and morphology. Spines were classified as stubby, thin, mushroom, and filopodia, according to previous work^20, 21^. 4-5 mice per genotype, 6 neurons per mouse, and 3 dendrites per neuron were analyzed.

### Hippocampus-dependent behavior

#### Novel location recognition

Mice were habituated to the open-field arena for 6 min, following exposure to two different objects in fixed positions for three consecutive trials of 6 min each. On day 2, mice explored the same arena with one object displaced to a novel position. Time spent exploring each object was manually scored in a blinded fashion to assess the discrimination index for the novel location.

#### Contextual fear-conditioning

Mice were trained using a 2-shock contextual fear conditioning paradigm, as previously described^4^. Mice received two periods of 30 s consisting of a paired cue light and a tone of 1,000-Hz, followed by a light foot-shock (2 s, 0.5 mA) separated by a 180-s interval (light foot-shock; Ugo Basile). Twenty-four hours later, mice were re-exposed to the same context for 3 min, and freezing levels (contextual) were measured using EthoVision XT system software (v14.0.1319, Noldus).

#### Barnes maze

Mice were tested on a large circular Barnes maze in which mice navigate using visual cues to an alternating escape hole over four trials on each day of the task, as described^4^. Search strategy classification was manually performed based on methods adapted from previous work^22^, categorized as localized, serial, random, scanning, focal, focal missense, targeted and direct. Each related strategy was categorized into non-hippocampus-dependent strategies (localized, serial, random, and scanning), and hippocampus-dependent strategies (focal, focal missense, targeted and direct). Cognitive performance of each trial on day 3 was scored according to the following, with a separation of 1 point between non-hippocampus-dependent and hippocampus-dependent strategies to account for additional cognitive complexity: localized=0, serial=1, random=2, scanning=3, focal=5, focal missense=5, targeted=6, direct=6. Focal and focal missense strategies are assigned “5”, as both are goal-directed strategies used without achieving the final target. Target and direct strategies were assigned “6” since both are goal-directed strategies with successful achievement of the final target.

#### Statistical analysis

Statistical analyses were performed using GraphPad Prism version 9.0 software (GraphPad Software) using tests described in text and figure legends, including student’s *t* test for 2-group comparisons, nested *t* tests with multi-level variables for 2-group comparisons, or chi-square analysis for 2-group comparisons of proportions. All experiments were performed in a randomized and blinded fashion.

## Results

### TIMP2 is expressed in hippocampal neurons, and its deletion induces transcriptomic changes

TIMP2 exhibits high expression in the hippocampus relative to other brain regions^4^. To identify the cellular sources of TIMP2 in the hippocampus and to begin to probe its function in this region, we examined TIMP2 protein levels by confocal microscopy in various subfields of the hippocampus of 2-month-old wildtype (WT) male and female mice. We found that TIMP2 is expressed in the hilus of dentate gyrus (DG), as well as CA3 and CA1 subfields of the hippocampus, and we found that the majority of TIMP2-expressing cells also stain for pan-neuronal marker, NeuN, arguing that the major cellular source of TIMP2 expression in these areas was neuronal (**Fig. 1A, 1B**). We did not detect differential TIMP2 neuronal expression in the hilus/DG, CA3, and CA1 subfields between males and females (**Supplementary Fig. 1A-C**).

**Figure 1.**
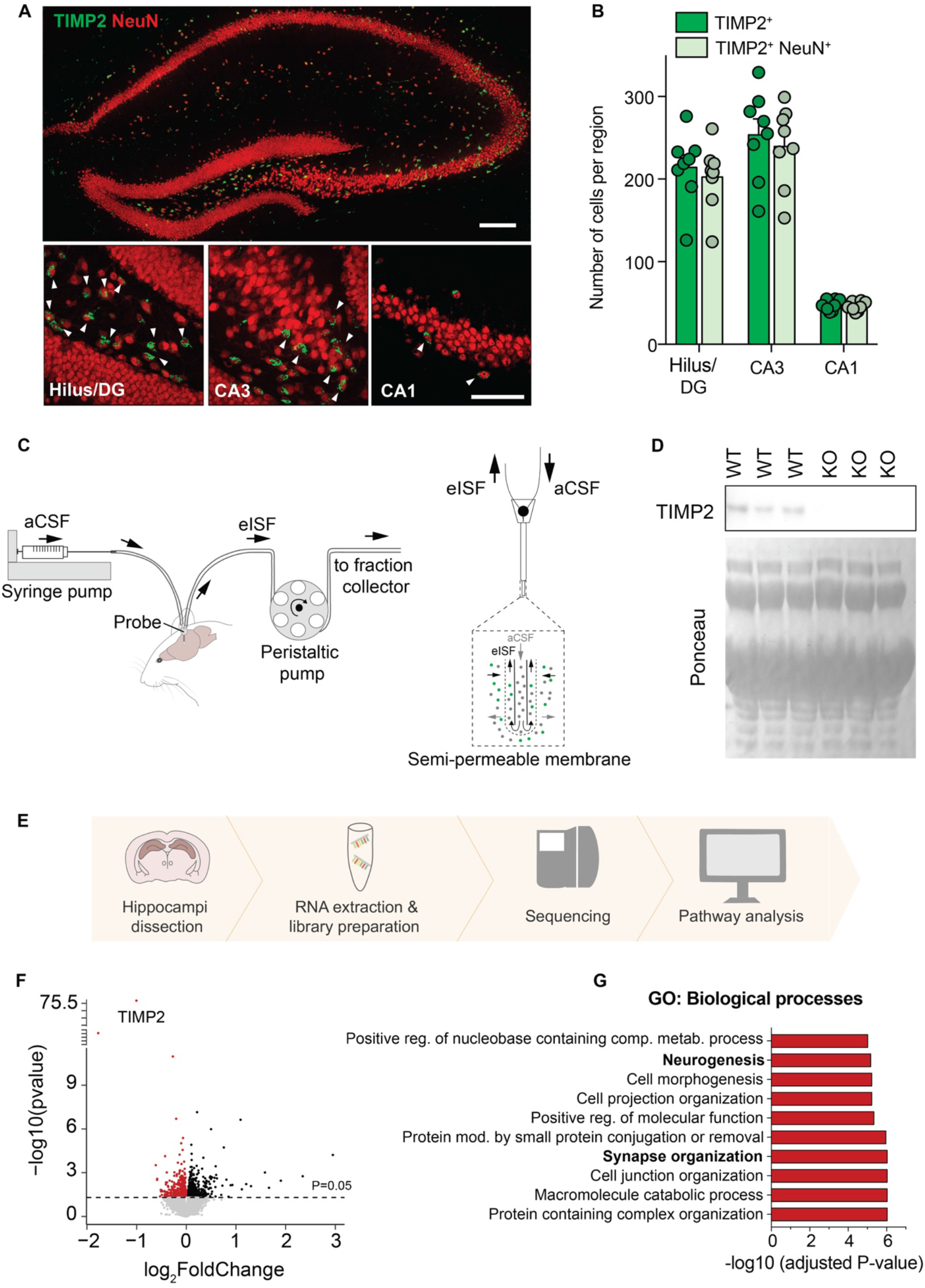
**TIMP2 is expressed in neurons of the hippocampus, and its deletion induces transcriptomic changes**. **(A)** Low-magnification view (upper image) of mouse hippocampus and high-magnification view (lower images) of hilus/DG, CA3, and CA1 subregions showing TIMP2^+^ cells and TIMP2^+^ cells staining for pan-neuronal nuclei marker NeuN. Scale bars, 200 μm and 20 μm (inset). **(B)** Quantification of the total number of TIMP2^+^ cells and TIMP2^+^ NeuN^+^ cells across hippocampal subregions in WT mice (2 months of age; N = 8, males and females). **(C)** Schematic representation of high molecular weight cut-off (1-MDa) *in vivo* microdialysis for assessing TIMP2 levels in mouse hippocampal ISF. **(D)** TIMP2 immunoblotting of hippocampal ISF dialyzed from 2-month-old WT and TIMP2 KO mice, with corresponding Ponceau S stain. **(E)** Schematic representation of bulk RNA-seq workflow performed in isolated WT and TIMP2 KO hippocampi (N = 13-17/group, sex-matched) for transcriptomic analysis. **(F)** Volcano plot showing the fold-change of genes (log2 scale) differentially expressed in hippocampus of TIMP2 KO vs. WT mice. Downregulated DEGs at *P* < 0.05 are highlighted in red (upregulated in black). **(G)** Top 10 significant pathways for downregulated DEGs from Gene Set Enrichment Analysis. Data are represented as mean ± SEM. DG, dentate gyrus; eISF, exchangeable interstitial fluid; aCSF, artificial cerebrospinal fluid.

Given the high level of TIMP2 expression seen in neurons in the hippocampus, we asked whether levels were elevated in the extracellular space adjacent to this cellular source. To measure extracellular levels of TIMP2 in this region, we implanted high molecular-weight cut-off (1-MDa) probes into hippocampi of freely moving, 2-3-month-old WT mice to measure interstitial fluid (ISF) levels of TIMP2 by *in vivo* microdialysis. TIMP2 was readily detectable in the ISF in hippocampus (**Fig. 1D**), and extrapolated values of exchangeable ISF TIMP2 were ∼10.1 ± 0.16 ng/mL (N=3), as measured at a flow rate of 1 μl/min. Importantly, the 21- to 22-kDa TIMP2 band was absent in the ISF of hippocampus from mice in which TIMP2 has been deleted (KO) (**Fig. 1D**).

The high levels of TIMP2 within neurons and the extracellular space of the hippocampus argues for unappreciated roles for its function locally in this region, prompting us to examine the cellular and molecular processes regulated by TIMP2. In order to identify putative pathways involved in TIMP2 function in the hippocampus, we isolated hippocampi from WT and TIMP2 KO male and female mice and processed the tissue for RNA-sequencing (**Fig. 1E**). A total of 905 differentially expressed genes (DEGs; *P<0.05*) were identified in TIMP2 KO compared to WT (**Fig. 1F**), with TIMP2 being the top DEG, as expected. Of these genes, 449 DEGs were downregulated in TIMP2 KO hippocampi (**Fig. 1F**). To gain insights into how the altered transcriptome may reflect altered biological processes, we performed gene set enrichment analysis (GSEA). Downregulated DEGs in TIMP2 KO were enriched for a number of Gene Ontology (GO) biological processes related to plasticity, including “cell morphogenesis”, “neurogenesis”, “cell projection organization”, and “synapse organization” (**Fig. 1G**). Upregulated DEGs in TIMP2 KO were enriched in functions that included “regulation of cell death and cell cycle”, among others (**Supplementary Fig. 1E**). Together, our data suggest that TIMP2 in the normal hippocampus may be involved in processes of plasticity.

### TIMP2 is necessary for adult neurogenesis in the normal dentate gyrus

To evaluate the extent to which our discovery-based pathway approach from RNAseq highlighted processes that may indeed be regulated by TIMP2, we examined the key cellular processes implicated. As one of the top downregulated pathways in hippocampus of TIMP2 KO vs. WT mice was “neurogenesis”, we used confocal microscopy to begin to address whether adult neurogenesis processes within the DG are indeed altered by absence of TIMP2. To examine whether TIMP2 regulates overall cell proliferation, we injected WT and TIMP2 KO mice with the proliferation marker bromodeoxyuridine (BrdU), which labels the pool of proliferating cells in the subgranular zone (SGZ) of DG (**Fig. 2A**). We found that the DG of TIMP2 KO mice had significantly fewer proliferating cells than in WT mice (**Fig. 2C**), as quantified by the total number of BrdU^+^ cells (**Fig. 2D**). We observed a similar result when analyzing the pool of Ki67^+^ cells in the SGZ (**Fig. 2E-F**), which marks a larger pool of proliferating cells in other phases. We next analyzed the pool of neural progenitor cells and the immature neuroblasts they ultimately give rise to, using SRY-Box Transcriptional Factor 2 (Sox2) and doublecortin (DCX) as markers, respectively. Deletion of TIMP2 resulted in a significant decrease in the number of Sox2^+^ neural progenitor cells (**Fig. 2G-H**), and of DCX^+^ immature neurons (**Fig. 2I-J**).

**Figure 2.**
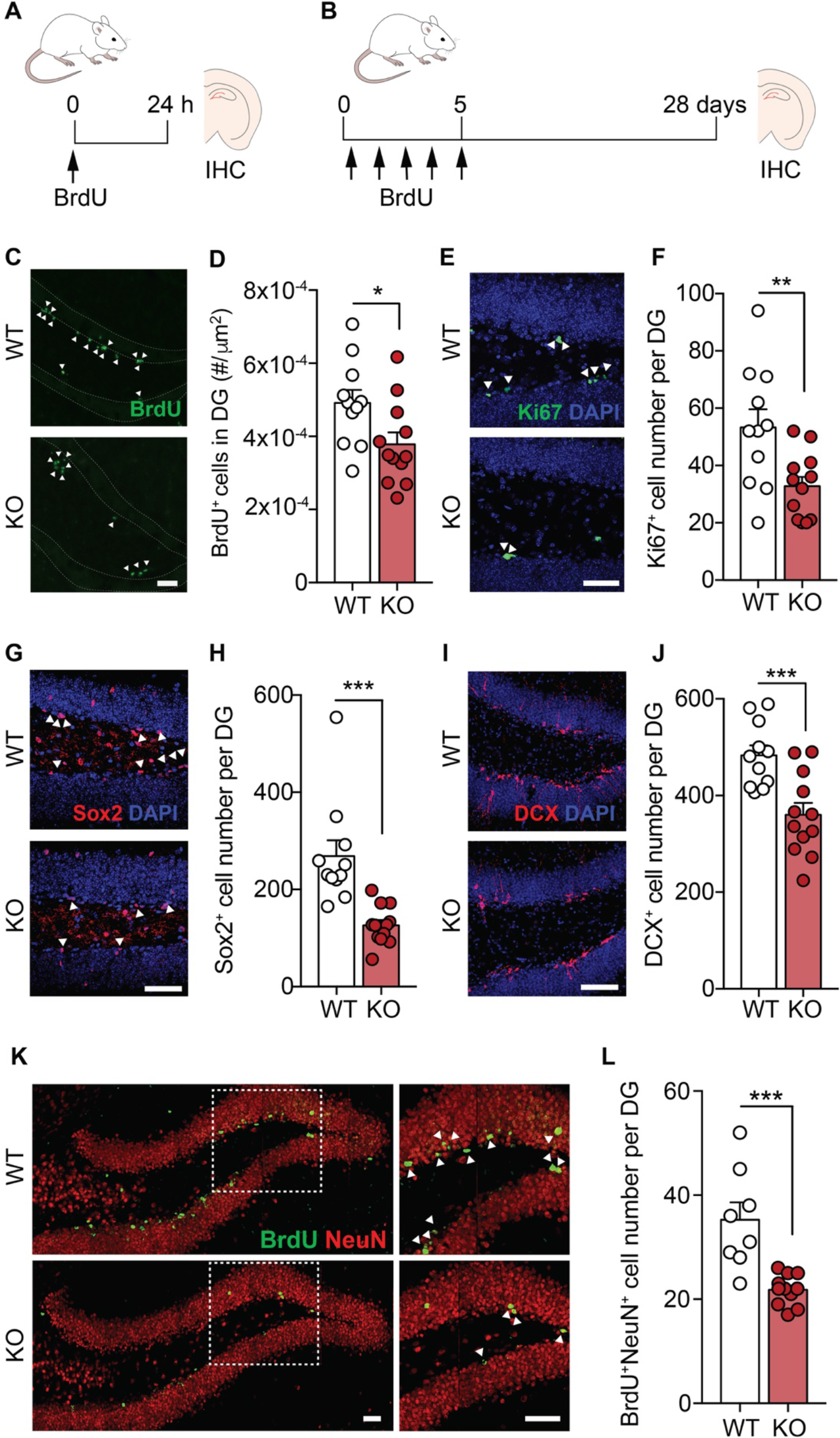
TIMP2 is necessary for adult neurogenesis in the normal dentate gyrus. (A) Schematic timeline of BrdU intraperitoneal injection protocol to label proliferating cells in the DG in isolated brain sections. **(B)** Schematic timeline of BrdU intraperitoneal injection protocol used for cell fate “survival” labeling in DG of isolated brain sections. **(C)** Representative confocal microscopy images of BrdU^+^ cells in the DG of WT and TIMP2 KO mice (2-3 months of age, N = 11-12 mice per group; arrowheads indicate BrdU^+^ cells; scale bar, 50 μm) with corresponding **(D)** quantification of number of BrdU^+^ cells in the dentate gyrus per unit area. **(E)** Representative confocal microscopy images of proliferating Ki67^+^ cells in the DG of WT and TIMP2 KO mice (2- 3 months of age, N = 11-12 mice per group; arrowheads indicate Ki67^+^ cells; scale bar, 50 μm) with corresponding **(F)** quantification of the number of Ki67^+^ proliferating cells per DG in WT and TIMP2 KO mice. **(G)** Representative confocal microscopy images of Sox2^+^ in the DG of WT and TIMP2 KO mice (2-3 months of age, N = 11-12 mice per group; arrowheads indicate Sox2^+^ cells; scale bar, 50 μm) with corresponding **(H)** quantification of Sox2^+^ neural progenitor cells in the DG of WT and TIMP2 KO mice. **(I)** Representative confocal microscopy images of DCX^+^ cells in the DG of WT and TIMP2 KO mice (2-3 months of age, N = 11-12 mice per group; scale bar, 50 μm) with corresponding **(J)** quantification of DCX^+^ immature neuroblasts in DG of WT and TIMP2 KO mice. **(K)** Representative confocal microscopy images of BrdU^+^ NeuN^+^ cells in DG of WT and TIMP2 KO mice (2-3 months of age. N = 8-11 mice per group; scale bar, 50 μm and 100 μm (inset)) with corresponding **(L)** quantification of newborn neurons (BrdU^+^ NeuN^+^ cells) in the DG of WT and TIMP2 KO mice. Data are represented as mean ± SEM. Student’s *t*-test for two-group comparisons. \**P*<0.05; \*\**P*<0.01; \*\*\**P*<0.001. Data points represent individual mice. IHC, immunohistochemistry; DG, dentate gyrus; DCX, doublecortin.

To examine whether this disrupted pool of neuroblasts corresponds to a differing number of surviving immature neurons, we injected TIMP2 KO and WT mice by pulse-chase labeling with BrdU for 5 days and quantified the number of label-retaining, surviving BrdU^+^ NeuN^+^ neurons in the SGZ of DG 4 weeks later (**Fig. 2B**). Using this paradigm, we found that the DG of TIMP2 KO mice had significantly fewer newborn neurons compared with the DG of WT mice (**Fig. 2K-L**). Together, these results are consistent with TIMP2-mediated roles in adult neurogenesis and related pathways uncovered by our transcriptomic analysis.

### TIMP2 regulates dendritic spine plasticity in the dentate gyrus

Our transcriptomic data in TIMP2 KO hippocampi suggest a role for TIMP2 acting within the hippocampus to modulate plasticity processes, including “synapse organization”, possibly at the structural level. To evaluate the impact of TIMP2 deletion on dendritic spines, the cellular substrates of memory storage^23^, we turned to a rigorous method to sensitively quantify dendritic spines and classify them. We performed Lucifer Yellow iontophoretic dye-filling of mature DG granule cells in isolated slices from TIMP2 KO and WT mice for high-resolution imaging and reconstruction (**Fig. 3A**)^21, 24^. We found that neurons in the DG of TIMP2 KO mice exhibited significantly reduced dendritic spine density compared to those in WT DG (**Fig. 3B, E**). We then used automated classification algorithms^21, 25^ to classify and quantify spines based on morphology, as a readout of spine plasticity. Intriguingly, we found a significantly higher proportion of immature thin spines in labeled DG neurons from TIMP2 KO mice compared to those in WT mice (**Fig. 3C, E**) and a significantly decreased proportion of mature mushroom spines (**Fig. 3D, E**) exhibited by TIMP2 KO relative to WT DG neurons. Mushroom spines are a particularly stable class of spines with lower turnover^26^ and are proposed to serve as structural correlates of memory^27^. Of note, our observations of spines in DG neurons of TIMP2 KO mice are in line with previous descriptions of altered synaptic function upon TIMP2 modulation^4^, including long-term potentiation, which was shown to be affected by modulating levels of TIMP2^4^. Collectively, these results suggest that TIMP2 may act to promote plasticity through regulation of adult neurogenesis and dendritic spine formation.

**Figure 3.**
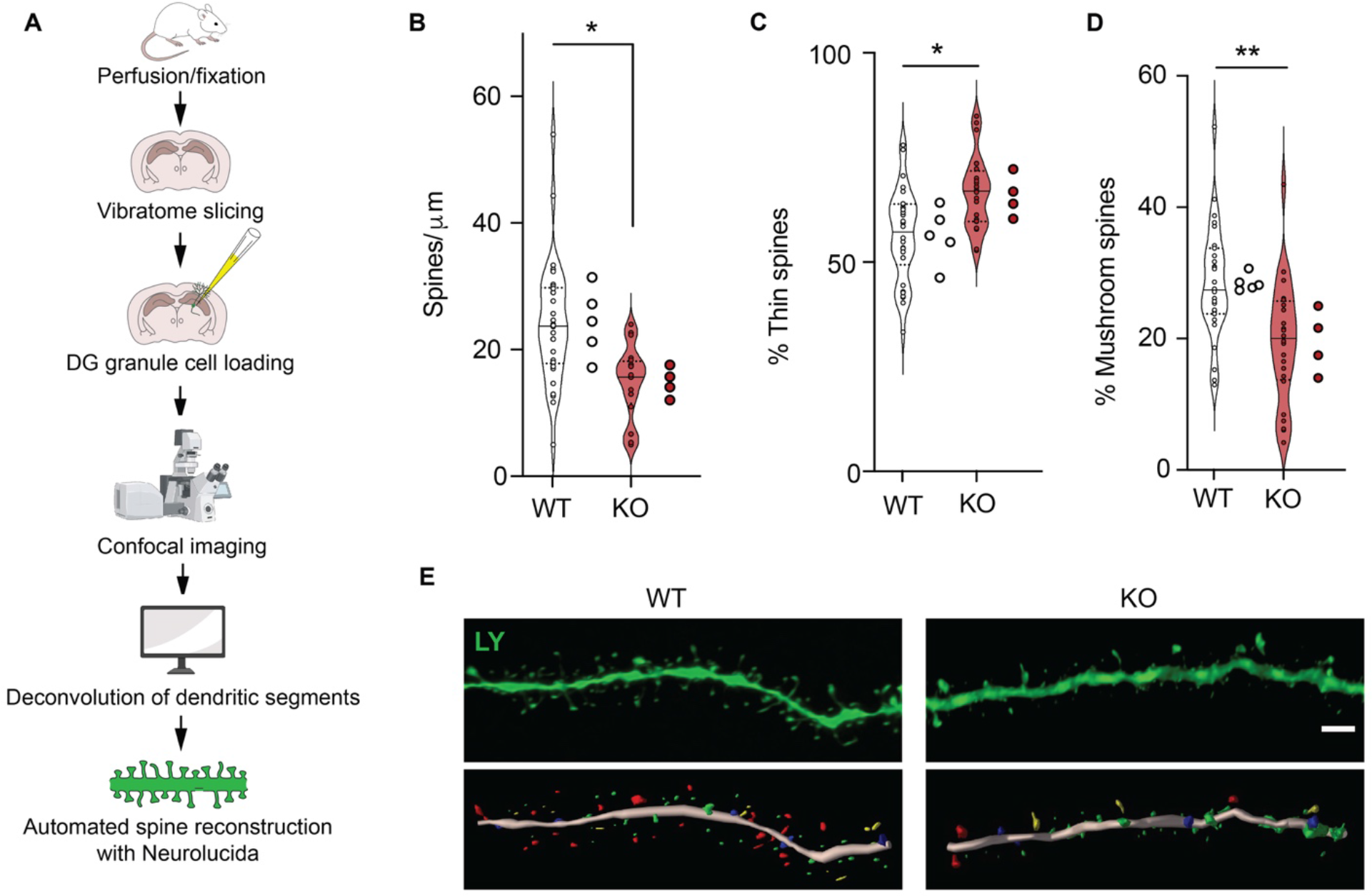
TIMP2 regulates dendritic spine plasticity within the dentate gyrus. (A) Schematic representation of the workflow for dendritic spine quantification in Lucifer Yellow-filled DG granule cells. **(B)** Overall dendritic spine density in DG granule cells iontophoretically labeled with Lucifer Yellow using sections isolated from six-month-old WT and TIMP2 KO mice (N = 6 neurons per mouse from N = 4-5 mice per group). **(C)** Quantification of the percentage of spines categorized according to “thin” spine heads or **(D)** “mushroom” spine heads. **(E)** Representative deconvolved confocal image of a dendritic segment from WT and TIMP2 KO Lucifer Yellow-labeled brain sections, and the downstream 3D reconstructions, with dendritic segment shown in pink, thin spines in green, stubby in blue, mushroom in red, and filopodia in yellow. Scale bar, 2 μm. Data are represented as mean ± SEM. Nested t-test for comparisons with neuron and mouse as levels. \**P*<0.05; \*\**P*<0.01. Data points represent neurons (left) and mice (right) for each group. LY, Lucifer Yellow

### TIMP2 is required for hippocampus-dependent memory function

The hippocampus is critical for spatial memory and learning, likely through regulation of adult neurogenesis in the DG (reviewed in^28^), as well as through changes in synaptic structure^29, 30^. The changes we observed in both adult neurogenesis and dendritic spine plasticity in TIMP2 KO hippocampi prompted us to investigate whether TIMP2 is involved in hippocampus-dependent memory and learning. We first utilized the novel location recognition task (**Fig. 4A**), in which mice use spatial cues in the arenas to learn and associate the positions of objects in space. Twenty-four hours after training, we observed that TIMP2 KO mice exhibited significantly decreased preference for the training object when displaced, as indicated by the reduced discrimination index (**Fig. 4B**), thus suggesting a critical role of TIMP2 in spatial memory in healthy mice. We next employed contextual and cued fear-conditioning to evaluate whether the mice differ in the ability to discriminate the previously aversive training context or a new cue-driven context (**Fig. 4C**). Assessment of context discrimination revealed significantly reduced freezing behavior by TIMP2 KO mice, suggesting impaired hippocampus-associated contextual discrimination in the absence of TIMP2 (**Fig. 4D-E**), while amygdala-associated cued freezing levels were unchanged (**Supplementary Fig. 2A**). We next examined whether more complex hippocampus-dependent spatial memory was altered in a Barnes maze assay in which mice use a variety of strategies while navigating an illuminated maze surface to an escape hole using spatial cues. Strategies were divided into non-hippocampus-dependent and hippocampus-dependent with increasing levels of cognitive demand (**Fig. 4F**)^22^. Analysis of the strategies used to reach the escape hole revealed that TIMP2 KO mice delayed the switch from non-hippocampal-dependent to hippocampal-dependent strategies while performing the task (**Fig. 4G**). On day 3 of the Barnes maze, a significant lower proportion of TIMP2 KO mice used hippocampus-dependent strategies compared to WT mice (**Fig. 4H**). Further categorization of cognitive performance by scoring the different strategies according to level of complexity related to spatial learning^31^ showed that TIMP2 KO mice achieved significantly lower cognitive scores relative to WT mice (**Fig. 4I**), consistent with impaired hippocampus-dependent function in TIMP2 KO mice while performing in the Barnes maze.

**Figure 4.**
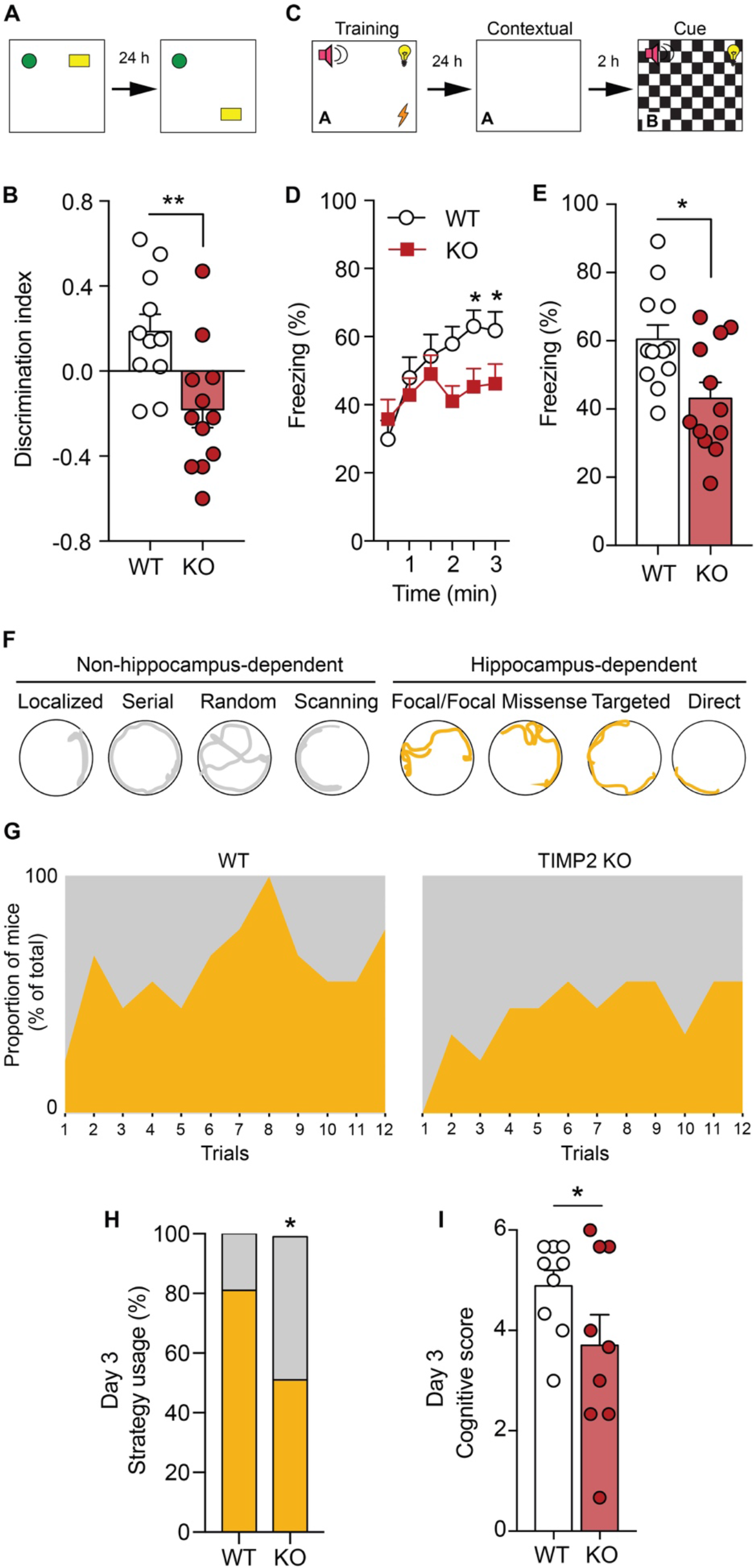
TIMP2 is required for hippocampus-dependent memory function. (A) Schematic of the novel location recognition assay. **(B)** Discrimination index for novel location recognition on day 2 for WT and TIMP2 KO mice (2-3 months of age, N = 11-12 mice per group). **(C)** Schematic of the contextual and cued fear-conditioning assay. **(D, E)** Freezing levels by interval and overall measured in the conditioned-fear context A in WT and TIMP2 KO mice (2-3 months of age, N = 11-12 mice per group). **(F)** Schematic diagram of modified Barnes maze with color coding for overall strategy classification. **(G)** Proportion of WT and TIMP2 KO mice using non-hippocampus-dependent (gray) and hippocampus-dependent (yellow) strategies during the testing trials (2-3 months of age, N = 9 mice per group). **(H)** Strategy utilization by WT and TIMP2 KO mice on day 3 of the Barnes Maze and corresponding **(I)** cognitive scores, ranked by strategy complexity for WT and TIMP2 KO mice on day 3 in the Barnes Maze. Data are represented as mean ± SEM. Student’s t-test for two-group comparisons in **(B, D, E)**, chi-square test in **(H)**, and nested t-test **(I)** for trial and mouse levels. \**P*<0.05, \*\**P*<0.01. Data points represent individual mice.

Importantly, we did not detect differences in motor function and overall activity in TIMP2 KO mice (**Supplementary Fig. 2B-J**). Motor coordination perturbations have been reported in TIMP2 mutant mice^13^, so we subjected the mice to a wide battery of motor function, coordination, and activity assays, including rotarod, open field, wire testing, pole testing, grip strength, none of which revealed changes in corresponding function between TIMP2 KO and WT mice (**Supplementary Fig. 2B-I**). We detected impaired clasping behavior in TIMP2 KO relative to WT mice (**Supplementary Fig. 2J**), as previously reported^13^, but this form of motor coordination does not impact performance observed for TIMP2 KO mice in the wide range of hippocampus-dependent tasks utilized. No differences in anxiety-related behaviors in the open field were observed between WT and TIMP2 KO mice (**Supplementary Fig. 2F**). Collectively, these results indicate a critical role for TIMP2 in regulation of hippocampus-dependent memory.

### TIMP2 ablation leads to accumulation of ECM in the hippocampus

We next investigated potential mechanisms that may account for how spine formation and adult DG neurogenesis are regulated by TIMP2. Outside the brain, TIMP2 has been characterized to be involved in ECM degradation and remodeling through its canonical binding partner MMP2^6, 7^. Brain ECM is dynamic, and accumulating evidence suggests a role for ECM and its components in the regulation of synaptic plasticity, including in spine remodeling^18, 32^, adult neurogenesis^33^, and cognition^33^. We first assessed the levels of pro-MMP2 in lysates of hippocampi from TIMP2 KO and WT mice. Interestingly, we found significantly elevated levels of MMP2, both at the gene (**Fig. 5A**) and protein (**Fig. 5B-C**) levels in the absence of TIMP2, reflecting an accumulation in response to loss in TIMP2 signaling and pointing to a potential disruption in ECM remodeling and turnover in hippocampus.

**Figure 5.**
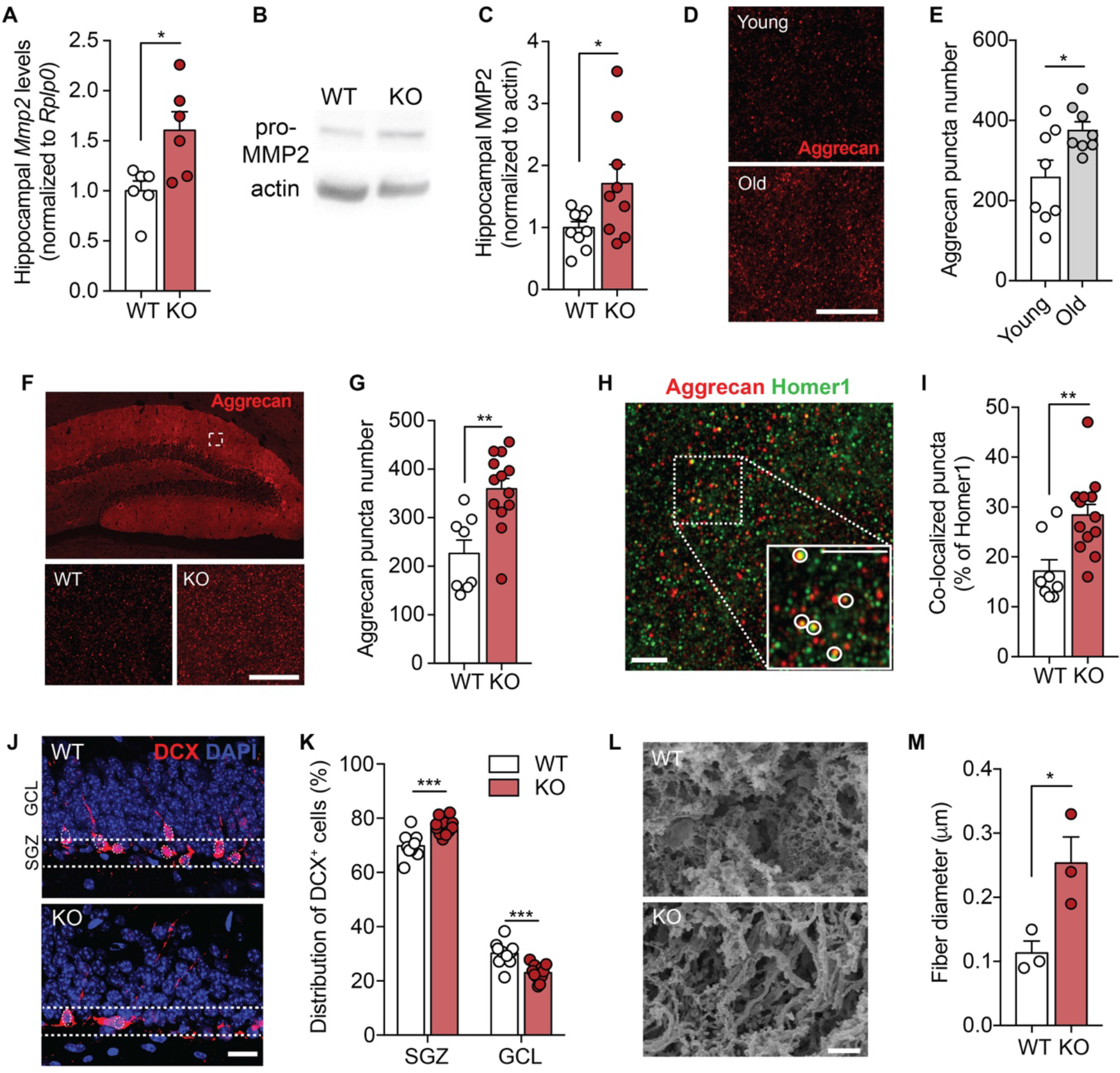
TIMP2 ablation leads to accumulation of ECM within the hippocampus. (A) *Mmp2* mRNA levels by qPCR from isolated hippocampus of WT and TIMP2 KO mice (2-3 months of age, N = 6 mice per group). **(B)** Immunoblotting of MMP2 from hippocampal lysates of WT and TIMP2 KO mice (2-3 months of age, N = 9 mice per group) with corresponding **(C)** quantification of MMP2 protein levels. **(D)** Representative confocal microscopy images of aggrecan puncta in the molecular layer of the DG from young (3-month-old) and old (21-month- old) mice (N = 8 mice per group; scale bar, 5 μm), with corresponding **(E)** quantification of aggrecan puncta number. **(F)** Representative confocal microscopy images showing aggrecan puncta in the molecular layer of DG at low-magnification (top) and high-magnification (bottom) of analysis inset from WT and TIMP2 KO mice (2-3 months of age, N = 8-13 mice per group; scale bar, 5 μm) with corresponding **(G)** quantification of puncta. **(H)** Representative confocal microscopy image indicating colocalized puncta (see inset) with corresponding **(I)** quantification of aggrecan and homer1 co-localization in the molecular layer of DG from WT and TIMP2 KO mice (2-3 months of age, N = 8-13 mice per group; scale bar, 2.5 μm). **(J)** Representative images showing the migration of DCX^+^ cells in the SGZ and GCL of WT and TIMP2 KO mice (2-3 months of age, N = 11-12 mice per group, scale bar, 20 μm with corresponding **(K)** quantification of DCX^+^ cells distributed in the subgranular zone (SGZ) and granule cell layer (GCL) of DG. **(L)** Representative surface scanning electron microscopy (SEM) from DG of WT and TIMP2 KO mice (2-3 months of age, N = 3 mice per group, scale bar, 2 μm, with corresponding **(M)** quantification of ECM microarchitecture in terms of fiber diameter measurements. Data are represented as mean ± SEM. Student’s t-test for two-group comparisons. \**P*<0.05, ** *P*<0.01, \*\*\**P*<0.001. Data points represent individual mice.

Recent work indicates that brain ECM levels increase with age in parallel with age-dependent cognitive decline^34^. In the brain, chondroitin sulfate proteoglycans (CSPGs) are the major class of ECM proteins^35, 36^. The proteoglycan aggrecan, a core CSPG expressed in perineuronal nets in the brain, is highly enriched in the ECM of the DG^37^ and has been linked to synaptic plasticity^38^ and memory^39, 40^. Using super-resolution confocal imaging, we find that aggrecan is significantly increased in the hippocampus of aged mice (**Fig. 5 D-E**), a change that occurs alongside the progressive decrease in TIMP2 expression^4^. We therefore analyzed aggrecan distribution in the hippocampus of young WT and TIMP2 KO mice and found that TIMP2 KO mice exhibited a significant increase in the density of aggrecan puncta in the DG molecular layer (**Fig. 5D-E**), the same area where we observed impaired spine remodeling in TIMP2 KO relative to WT mice (**Fig. 3**). To further determine if TIMP2 remodeling of the ECM could affect accumulation of ECM near synapses given the synaptic deficits we observed in these neurons, we analyzed aggrecan colocalization with the postsynaptic protein Homer1. We observed that the DG of TIMP2 KO mice exhibited significantly increased colocalization of aggrecan with Homer1 relative to that observed in WT DG (**Fig. 5F-G**). These data indicate that TIMP2 normally promotes ECM remodeling and limits its deposition near synapses.

We next further characterized the relationship between TIMP2’s effect on ECM and adult neurogenesis, considering the known roles of ECM and its related components in the regulation of proliferation and migration of newborn neurons (reviewed in^41^). To explore this possibility, we evaluated the migration of newborn neurons toward the molecular layer by examining the distribution of DCX^+^ immature neuroblasts in the SGZ relative to that in the GCL of the DG. This analysis revealed significantly decreased distribution of DCX^+^ immature neurons in the GCL of TIMP2 KO mice (**Fig. 5H-I**), reflecting a process whereby migration of newborn neurons within the DG is apparently impaired in the absence of TIMP2. These data suggest that the absence of TIMP2 leads to a tighter ECM structural environment that acts to spatially restrict the migration of newborn neurons and plasticity of synapses, thus affecting overall adult neurogenesis and spine dynamics. In support of this concept, we measured structural changes in the ECM of TIMP2 KO mice by performing surface scanning electron microscopy (SEM) on decellularized brains isolated from TIMP2 KO and WT mice. Interestingly, we detected altered ECM microarchitecture in the form of thicker ECM fibers (**Fig. 5J-K**) in hippocampi from TIMP2 KO compared to WT mice. Together, our results suggest that TIMP2 modulates the flexibility of the ECM within the hippocampus to regulate plasticity processes, including adult neurogenesis and spine plasticity.

### Neuronal TIMP2 regulation of the extracellular matrix regulates hippocampus-dependent cognitive function and adult neurogenesis

To gain further mechanistic insight into TIMP2’s role in the function of the hippocampus with increased cellular resolution, we generated *Timp2*-floxed (TIMP2^fl/fl^) mice using two-cell homologous recombination (2C-HR)-CRISPR/Cas9-based genome editing. We inserted two *loxP* sites flanking exon 2 of the *Timp2* gene transcript (**Supplementary Fig. 3A-B**) and then crossbred *Timp2*^fl/fl^ mice with Syn^Cre/+^ mice (**Supplementary Fig. 3C**) to specifically excise *Timp2* in neurons. Immunoblotting against TIMP2 from isolated hippocampus lysates of Syn^Cre/+^; TIMP2^fl/fl^ or TIMP2^fl/fl^ control mice confirmed that conditional targeting of TIMP2 resulted in significantly reduced TIMP2 levels compared to TIMP2^fl/fl^ controls in hippocampus (**Supplementary Fig. 3D-E**). We performed further validation by performing confocal microscopy in hippocampus of Syn^Cre/+^; TIMP2^fl/fl^ or TIMP2^fl/fl^ control mice to examine the cells in hippocampus expressing TIMP2. Quantification of TIMP2^+^ NeuN^+^ cells in the hilus/DG, CA3 and CA1 regions revealed significant abrogation of TIMP2 in neurons upon its conditional deletion (**Supplementary Fig. 3F-G**). These data validate our model as a tool for addressing the role of neuronal TIMP2 in regulating the key cellular and behavioral phenotypes observed in the context of global TIMP2 deletion.

Given that we found that TIMP2 was expressed by neurons in DG and present at significant levels in the extracellular space adjacent to these cells, we sought to determine whether the pool of neuronal TIMP2 is specifically involved in hippocampus-dependent memory. Using the Syn^Cre/+^; TIMP2^fl/fl^ mice we generated and validated (**Supplementary Fig. 3**), we created cohorts of Syn^Cre/+^; TIMP2^fl/fl^ mice and TIMP2^fl/fl^ control littermates to evaluate performance in spatial memory tasks. We found that Syn^Cre/+^; TIMP2^fl/fl^ mice exhibited significantly decreased preference for the displaced object 24 hours after training in the testing phase compared to TIMP2^fl/fl^ controls, as indicated by a reduced discrimination index in these mice (**Fig. 6A**). Further assessment in the contextual portion of fear-conditioning revealed a significant reduction in freezing levels by Syn^Cre/+^; TIMP2^fl/fl^ mice compared to TIMP2^fl/fl^ controls, suggesting impaired contextual memory in the absence of neuronal TIMP2 (**Fig. 6B-C**). Cued freezing levels were indistinguishable between the groups of mice (**Supplementary Fig. 4A**). We then subjected the mice to Barnes maze testing to examine performance in search strategies to find the escape hole. Our search strategy analysis showed that a smaller proportion of Syn^Cre/+^; TIMP2^fl/fl^ mice used hippocampus-dependent strategies compared to TIMP2^fl/fl^ mice (**Fig. 6D-E**). Scoring their search strategy utilization by complexity shows that Syn^Cre/+^; TIMP2^fl/fl^ mice score lower than their TIMP2^fl/fl^ counterparts (**Fig. 6F**), suggesting TIMP2 from neurons is required for normal hippocampus-related memory and learning. Importantly, we did not detect differences in overall motor function in terms of coordination (**Supplementary Fig. 4B-C**), overall activity in the open field (**Supplementary Fig. 4D-E**), grip strength or motor learning (**Supplementary Fig. 4G-I**) and clasping behavior (**Figure S3J**) in Syn^Cre/+^; TIMP2^fl/fl^ mice relative to TIMP2^fl/fl^ mice. No differences in anxiety-related behaviors in the open field were observed between TIMP2^fl/fl^ and Syn^Cre/+^; TIMP2^fl/fl^ mice (**Supplementary Fig. 4F**).

**Figure 6.**
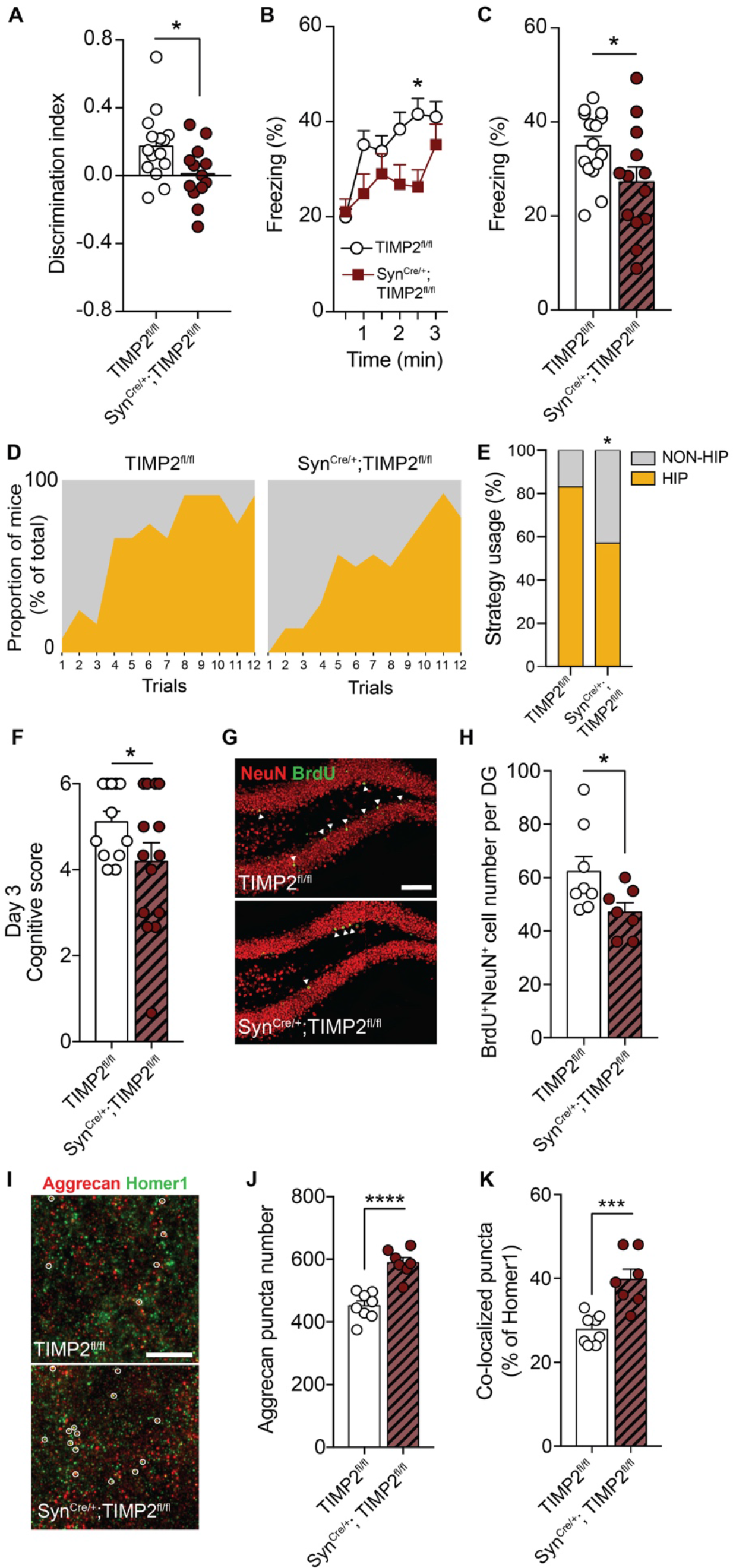
Neuronal TIMP2 regulation of the extracellular matrix regulates hippocampus- dependent cognitive function and adult neurogenesis. (A) Discrimination index for novel location recognition on day 2 and (B, C) contextual fear- conditioning freezing levels in TIMP2^fl/fl^ and Syn^Cre/+^; TIMP2^fl/fl^ mice (2-3 months of age, N = 13- 15 mice per group). (D) Proportion of mice of each genotype using non-hippocampus-dependent (gray) and hippocampus-dependent (yellow) search strategies during the testing trials of TIMP2^fl/fl^ and Syn^Cre/+^; TIMP2^fl/fl^ mice in Barnes maze (2-3 months of age, N = 12-14 mice per group) with (E) quantification of the percentage of mice using these strategies by on day 3. (F) Cognitive complexity scores for TIMP2^fl/fl^ and Syn^Cre/+^; TIMP2^fl/fl^ mice based on strategies used on day 3 of Barnes maze. (G) Representative confocal microscopy images of BrdU^+^ and NeuN^+^ cells in DG of TIMP2^fl/fl^ and Syn^Cre/+^; TIMP2^fl/fl^ mice (2-3 months of age, N = 7–8 mice per group, scale bar, 100 μm) with corresponding (H) quantification of BrdU^+^ NeuN^+^ newborn neurons in the DG. (I) Representative confocal microscopy images of aggrecan and homer1 puncta in molecular layer of DG of TIMP2^fl/fl^ and Syn^Cre/+^; TIMP2^fl/fl^ mice (N = 7-8 mice per group, scale bar, 5 μm) with corresponding (J) quantification of aggrecan puncta and (K) co-localized puncta of aggrecan and homer1, as indicated by overlaid circles. Data are represented as mean ± SEM. Student’s *t*-test for two-group comparisons in (A-C), (H), and (J, K), chi-square test in (E), and nested t-test (F) with trial and mice as levels. \**P*<0.05, ** *P*<0.01, \*\*\**P*<0.001, \*\*\*\**P*<0.0001. Data points represent individual mice.

To test whether the local pool of neuronal TIMP2 affects the survival of newly generated neurons, we analyzed the pool of BrdU^+^ NeuN^+^ cells after pulse-chase labeling with BrdU, as described earlier (**Fig. 2B**). Indeed, we observed that the deletion of TIMP2 within neurons resulted in a significant decrease in the number of BrdU^+^ NeuN^+^ newborn neurons compared to TIMP2^fl/fl^ controls (**Fig. 6G-H**). To probe potential mechanisms by which TIMP2 expressed locally from neurons affects hippocampus-dependent memory and adult neurogenesis, we next addressed ECM-related remodeling in the DG of Syn^Cre/+^; TIMP2^fl/fl^ mice. TIMP2 deletion in neurons in Syn^Cre/+^; TIMP2^fl/fl^ mice resulted in a significant increase in the density of aggrecan-positive puncta compared to TIMP2^fl/fl^ controls (**Fig. 6I-J**), as well as increased colocalization of aggrecan with Homer1 (**Fig. 6I, 6K**), consistent with an accumulation of ECM components adjacent to synapses, as observed earlier. Together, our results show that the pool of neuronal TIMP2 is necessary to promote ECM remodeling to enable synaptic plasticity and adult neurogenesis to facilitate normal hippocampus-dependent memory.

## Discussion

Our study illustrates how alterations in the ECM within the hippocampus by TIMP2 can directly regulate features of plasticity, including dendritic spine plasticity, adult neurogenesis, and hippocampus-dependent memory. We find that TIMP2 is highly expressed in neurons within the hippocampus and found at high levels in the brain extracellular space. Its removal induces structural synaptic changes and deficits in adult neurogenesis, as predicted from changes observed at the transcriptomic level, both of which result in impairments in hippocampus-dependent memory function. We demonstrate that TIMP2’s role in ECM remodeling is associated with changes in the flexibility of synaptic plasticity, impinging on processes like neuronal migration and differentiation. We generate and characterize a novel mouse model for conditional deletion of TIMP2 within neurons, highlighting molecular mechanisms through which neuronal TIMP2 promotes synaptic plasticity through regulation in the ECM to facilitate normal memory.

The development and maintenance of synapses is critical in the establishment of functional neuronal circuits, and emerging evidence indicates a role for the ECM and its components in regulating synapse structure and morphology^41–43^. We used high-resolution imaging of labeled granule cells in the intact hippocampus of TIMP2 KO and WT mice to show that TIMP2 regulates dendritic spine density, with a loss in KO mice in the class of spines involved in memory, mushroom spines^26^. Moreover, when TIMP2 is targeted either globally or specifically within neurons, ECM components like aggrecan accumulate around hippocampal synapses, and the ECM is more dense, as shown by our SEM data, suggesting that TIMP2 acts directly within the hippocampus to modulate plasticity on a structural level. In agreement with its role in modulating plasticity, we also find that TIMP2 regulates the process of adult neurogenesis. While immature neuroblasts were unaltered in mice with systemic modulation of TIMP2^4^, presumably due to the window over which TIMP2 was modulated being too short to reflect changes in DCX^+^ cells, we now show that removal of TIMP2, both globally and from neurons in the hippocampus, results in disruption in neural progenitor proliferation and maturation. This is in line with *in vitro* evidence indicating that TIMP2 expression was significantly upregulated upon neuronal differentiation^44^, and that it may be involved in differentiation^12^. Intriguingly, we observed that TIMP2 deletion affected the migration of immature DCX^+^ neurons towards the molecular layer, arguing for a role of ECM in regulating neuronal migration^45^. It has been documented that ECM molecules participate in neurogenesis^41^, though the mechanisms by which they regulate neuronal differentiation are unclear. CSPGs are known to surround new neurons in the hippocampus and have recently been shown to be important in the production of new neurons^33^, and ECM-related proteins like reelin also play critical roles in neuronal migration^45^.

TIMP2 is traditionally recognized for its unique dual role in regulating both the activation of pro-MMP2 and the inhibition of MMP2 activity necessary for ECM remodeling^6, 7^. We observed that deletion of TIMP2 increased levels of *Mmp2* and pro-MMP2 levels within hippocampus, perhaps reflecting compensatory responses to reduced TIMP2 and impaired MMP2 activation, as described outside of the brain^6, 7^. Accumulated pro-MMP2 appears to translate to less remodeling of the ECM and its resulting accumulation around cells of the DG neurogenic niche we observe. A proposed mechanism of spine stabilization and remodeling by ECM includes a structural-spatial restriction mechanism in which ECM components, including CSPGs, form a matrix around dendritic spines to provide extracellular rigidity and physically restrict spine motion^46^. MMPs themselves can modulate structural plasticity by loosening the ECM structure, allowing for a more permissive environment^41^. Moreover, stiffness of the surrounding environment determined by ECM complexity has been implicated in regulating cell migration^47, 48^, including for neural stem cells *in vitro*^49–51^. This modulation has been proposed to be among fundamental cell fate-determining factors in neural development and neurological disorders^52–54^. Our results argue that ECM accumulation in the absence of TIMP2 leads to a denser network that creates physical restrictions that impact turnover of synapses from thin into mature mushroom-shaped spines, while impeding proliferation and migration related to adult neurogenesis.

In line with reports linking disrupted synaptic plasticity and neurogenesis with cognitive dysfunction^41^, we show that TIMP2-related changes in hippocampus plasticity create significant memory impairments. Both global and neuron-specific targeting of TIMP2 levels leads to deficits in hippocampus-dependent memory tasks. Our results strongly reinforce a surprising link among TIMP2, synaptic plasticity, and memory in the healthy hippocampus through regulating spine dynamics and neurogenesis. Importantly, the cognitive phenotypes described here cannot be attributed to TIMP2’s role in other brain regions related to motor function, though earlier work in younger mice described subtle deficits in coordination upon TIMP2 deletion^13^, primarily related to cerebellar alterations deletion^13^. We show that TIMP2-mediated roles in ECM to regulate cellular phenotypes are associated with changes in MMP2. TIMP2 likely interacts with many MMPs and other binding partners to mediate functional changes under different conditions, including alpha3beta1 integrin^12^. It is possible that TIMP2 exerts its diverse biological functions through interactions with different binding partners, or that it exerts its activities differently *in vivo* versus *in vitro*, all of which will need to be further explored in future studies. Moreover, while we mostly find TIMP2 expressed in neurons in our study, its expression in other cell types may be important in certain pathological contexts^55^ to be explored further, especially as a role for microglia in remodeling the ECM to promote synapse plasticity in the hippocampus has been recently described^18^.

Dysregulation of the ECM is implicated in cognitive dysfunction that occurs in aging^56^ and in disease^57^. The progressive decline in TIMP2 expression with age^4^ occurs in parallel with deficits in memory^58^ and ECM accumulation, as demonstrated here, while systemic treatment with TIMP2 revitalizes hippocampus-related function in aged mice^4^. Moreover, ECM clearance is beneficial in mouse models of Alzheimer’s disease pathology^59^. It has been increasingly appreciated that adult neurogenesis in the human DG is dysregulated in Alzheimer’s disease^60^. Defining the precise molecular mechanisms accounting for how modulators of the ECM regulate the healthy brain may lead to development of novel approaches for cognitive repair in contexts of aging or neurodegeneration.

## Acknowledgments

We gratefully acknowledge the contributions of the Mouse Genetics and Gene Targeting (MGGT) CoRE facility (Kevin Kelley, ISMMS), which is supported by ISMMS and the National Cancer Institute of the NIH under award P30CA196521, for two-cell stage embryo microinjections for generating TIMP2^fl/fl^ mice. We also thank Chingwen Yang of the CRISPR and Genome Editing Center (Rockefeller University) for assistance generating constructs to develop the TIMP2^fl/fl^ line. We thank Yihang Wang (ISMMS) for RNA isolation support, Nikos Tzavaras and Deanna Benson of the Microscopy CoRE and Advanced Bioimaging Center (ISMMS) for advice and equipment for Airyscan microscopy and image analysis, Lee Cohen-Gould of the Electron Microscopy & Histology Services of Weill Cornell Medicine Microscopy & Image Analysis Core for SEM sample preparation, Jorge Morales of the Microscopy Facility (CCNY) for SEM acquisition support, and Ron Gordon and Andy Leonard of the EM Core (ISMMS) for advice and sample preparation for SEM. This work was funded by the National Institute on Aging (R01AG061382 (J.M.C), RF1AG072300 (J.M.C.), T32AG049688 (B.M.H.), and R01AG061382-02S1 (J.M.C., S.M.P)).

## Author contributions

A.C.F. and J.M.C. conceived and designed the experiments, performed data analysis and interpretation, and wrote the manuscript. A.C.F. conducted the experiments. B.M.H. performed and analyzed microdialysis experiments. S.M.P analyzed data for RNAseq experiments. H.L. and J.D.Z performed or analyzed motor or Barnes maze experiments. M.V. performed intracellular dye injections for spine labeling, and analyzed/interpreted data with P.R.H. T.K. and T.A. assisted with design and analysis of TIMP2^fl/fl^ validation data. All authors approved the final manuscript.

## Conflict of interests

J.M.C. is listed as a co-inventor on patents for treating aging-associated conditions, including the use of young plasma administration (US10688130B2) or youth-associated protein TIMP2 (US10617744B2), the latter of which is licensed to Alkahest, Inc.

Supplementary Materials and Methods and Supplementary Figures are available at the end of the document.

## Supplementary Information, Materials and Methods and Supplementary Figures

### Supplementary Materials and methods

#### Generation of TIMP2-floxed mice

To achieve conditional deletion of *Timp2*, a loxP-flanked *Timp2* allele (TIMP2^fl/fl^) was generated by the two-cell homologous recombination (2C-HR)-CRISPR/Cas9-based genome editing method, as previously described^1^. Briefly, two loxP sites were inserted into the *Timp2* gene flanking exon 2, with homologous arms (HA) on each side. Two point-mutations (C928->A near the end of the left arm and G1795->T at the 5’ junction of the right arm) were introduced to disrupt the PAM sequence to avoid re-cutting of the recombined allele. Three nucleotides adjacent to each loxP site were altered to create restriction sites for diagnostic purposes: XbaI near the upstream loxP, and HindIII near the downstream loxP site. The two-cell stage embryo microinjection procedure was performed as in^1^, using Cas9-mSA mRNA, biotinylated double-stranded PCR product (1016 nts), and pre-assembled ctRNA prepared accordingly^1^. F0 mice were screened by sequencing from TOPO cloned PCR product, using tail genomic DNA, amplified from the TIMP2 target region, and using the following primers: FW-AGCGACCGATAAGCAGGAAG, Rev-CACTAGCAGACACCACCACA. Identified founders were crossed with wild-type (WT) C57BL/6J mice, purchased from The Jackson Laboratory, to produce heterozygous mice.

To achieve neuron-specific *Timp2* deletion, neuron-specific heterozygous Synapsin (Syn)- Cre (Syn^Cre/+^) transgenic mice^2^ (JAX stock #003966) were crossbred with TIMP2^fl/fl^ mice to generate Syn^Cre/+^; TIMP2^fl/+^ progeny. The F1 mice were then crossed to generate Syn^Cre/+^; TIMP2^fl/fl^ and TIMP2^fl/fl^ littermate controls. Cre expression was maintained in the female mice for breeding^2^. All animals were maintained on a C57Bl/6J background.

#### Genotyping

Genotyping of TIMP2^fl/fl^ mice was performed using the following primers: *Primer 1*: 5’- TGACCCCTCTGCTCTAGTCC, *Primer 2:* 5’-GCTCCGTCTCTTCGTCCATC (**Figure 6A**).

Genomic DNA was extracted from the tail using the QIAamp Fast DNA Tissue Kit (Qiagen). PCR was performed in a total volume of 50 µl at 94°C for 2 min, 94°C for 30 s, 61°C for 30 s, 68°C for 2 min and 15 sec for 10 cycles, then 94°C for 30 s, 56°C for 30 s, 68°C for 2 min and 15 s for 25 cycles, and ending at 72°C for 5 min using Prime STAR (#R050A, TaKaRa, Shiga, Japan). For loxP site verification based on restriction digest, the PCR product (2.1 kb) was digested with XbaI and HindIII restriction enzymes (New England Biolabs, Ipswich, MA, USA) at 37°C for 1 hour, followed by 10 min at 65°C for heat inactivation. Digested PCR products were run on an agarose gel and imaged for the identification of founders: HindIII (WT: 2133 bp; upstream loxP positive: 829 bp and 1,304 bp); Xba1 (WT: 2133 bp; downstream loxP positive: 566 bp and 1,567 bp).

#### In vivo microdialysis

For guide cannula implantation into the hippocampus, mice were anesthetized with 2% isoflurane and head was shaved prior to being fixed in a stereotaxic apparatus (Stoelting Co., Wood Dale, IL, USA). After preparing skin with 70% ethanol and povidone-iodine, an anterior-to-posterior incision was made along the midline of the head to expose the skull. Skull position was leveled to within 0.1 mm along the bregma-lambda axis, and at 2.2 mm to the left and right of midline on the lateral axis. Holes were drilled to target the left hippocampus using the stereotaxic coordinates: bregma -3.1 mm, 2.5 mm lateral. A second hole was made diagonally to the first, to serve as an anchoring position with a bone screw. To target caudal hippocampus, AtomosLM Guide Cannula (PEG-12, Eicom, London, UK) was inserted at a 12° angle, 1.2 mm below the dura mater. Dental cement was applied to secure the cannula, and the skin was secured using a surgical adhesive glue. An AtmosLM Dummy Cannula (PED-12, Eicom) was inserted into the guide cannula and secured with a plastic cap nut. Animals were then placed in a clean cage on a heating pad and allowed to recover from surgery.

Approximately 12 hours following surgery, mice underwent 1000-kDa *in vivo* microdialysis. The inlet port of AtmosLM Microdialysis Probe (PEP-12-02, Emicon) was connected to a syringe pump (kdScientific, Holliston, MA, USA) perfusing artificial cerebrospinal fluid (aCSF), in mM: 1.3 CaCl2, 1.2 MgSO4, 3 KCl, 0.4 KH2PO4, 25 NaHCO3, 122 NaCl, pH 7.35, at a rate of 1.2 μl/min^3^. The outlet of the probe was connected to the peristaltic pump (MAB 20, SciPro, New York, NY, USA), which was calibrated to obtain a pull rate between 1-1.1 μl/min. After the probe was connected to the push-pull mechanism, the mice were briefly anesthetized with isoflurane to allow for the removal of the dummy cannula and insertion of the probe into the hippocampus. The probe was secured with a cap nut, and a plastic collar was loosely placed around the neck of the mouse. Mice were then moved to a Raturn (Stand-Alone Raturn System, BASi, West Lafayette, IN, USA) and tethered by the collar to avoid tangling microdialysis tubing (FEP tubing 0.65 mm OD x 0.12 mm ID, BASi), while allowing for free movement of the mouse during the sample collection period. Samples were collected hourly in a refrigerated fraction collector (MAB 85 Fraction Collector, SciPro) and frozen at −80°C after collection. The procedure was performed under constant light conditions, and food and water were provided *ad libitum*.

#### Bulk RNA-sequencing

RNA-sequencing was performed on hippocampi dissected from WT and TIMP2 KO male and female mice. Dissected hippocampi were preserved in RNAlater (Invitrogen) overnight at 4°C before storing at −80°C until use. RNA was extracted using the RNeasy Mini Kit (Qiagen), according to the manufacturer’s instructions. RNA quality was measured using the Agilent TapeStation Bioanalyzer (Agilent Technologies), and all samples exhibited RNA Integrity Number (RIN) > 8. cDNA libraries were prepared with poly(A) selection and sequenced using Illumina Hiseq (2x150bp paired-end) (Genewiz). At least 25 M clean reads were generated from each sample. The reads were mapped to the *Mus musculus* GRCm38 reference genome available on ENSEMBL, using STAR aligner (v.2.5.2b). After extraction of gene hit counts, the gene hit counts table was used for downstream differential expression analysis, using DESeq2 software. The Wald test was used to generate *P*-values and log2 fold changes. Differentially expressed genes (DEGs; *P*<0.05) were used for Gene Set Enrichment Analysis (GSEA; https://www.gsea-msigdb.org/gsea/index.jsp). Volcano plot was generated with R v4.1.2.

#### Immunohistochemistry and microscopy

For all immunohistochemistry experiments, mice were anesthetized with a cocktail of ketamine (90 mg/kg) and xylazine (10 mg/kg) and transcardially perfused with ice-cold 0.9% saline. Brains were dissected, and hemibrains isolated for immunohistochemistry were postfixed in 4% paraformaldehyde (PFA) for 48 hours at 4°C before preservation in 30% sucrose (Sigma, St. Louis, MO, USA) solution in phosphate-buffered saline (PBS). Hemibrains were sectioned coronally at 40-µm thickness on a freezing-sliding microtome (SM2010R, Leica, Deer Park, IL, USA), and sections were stored in cryoprotective medium at −20°C.

For immunohistochemistry, free-floating brain sections were permeabilized with Tris-buffer saline (TBS) plus 0.05% Tween-20 (TBST) and incubated in a blocking solution consisting of 10% normal goat (#S-1000, Vector Laboratories, Newark, CA, USA) or donkey serum (#017- 000-121, Jackson ImmunoResearch, West Grove, PA, USA) in TBST. Sections were then incubated with primary antibodies in 2% goat or donkey serum in TBST overnight at 4°C. Sections were incubated with primary antibodies, including the following: TIMP2 (1:100; #AF971, R&D Systems, Minneapolis, MN, USA), NeuN (1:200; #MAB377, Millipore, Burlington, MA, USA), doublecortin (DCX; 1:200; #4604S, Cell Signaling, Danvers, MA, USA), Sox2 (1:200; #sc365823, Santa Cruz, Dallas, TX, USA), Ki67 (1:200; #14-5698-82, Thermo Fisher, Waltham, MA, USA), aggrecan (1:500; #AB1031, Millipore), Homer1 (1:500; #160006, Synaptic Systems, Göttingen, Germany). The next day, sections were washed in TBST three times for 5 min each and incubated with Alexa Fluor-conjugated secondary antibodies at 1:200 in TBST for 1 hour at RT. Secondary antibodies used were: Alexa Fluor 594-labeled donkey anti-rabbit (#A-21206), Alexa Fluor 594-labeled donkey anti-mouse (#A-21203), Alexa Fluor 488-donkey anti-rat (#A-21208), Alexa Fluor 488-donkey anti-goat (#A-11055), Alexa Fluor 647-donkey anti-mouse (#A-31571), Alexa Fluor 594-goat anti-rabbit (#A-11012), Alexa Fluor 5488-goat anti-chicken (#A-11039), all from Thermo Fisher. Brain sections were then washed, stained with 4′,6-diamidino-2-phenylindole (DAPI, Sigma) for 15 min to visualize nuclei, mounted and coverslipped using Prolong Gold (Life Technologies) and dried overnight before imaging. For BrdU staining, brain sections were pre-treated with 3M HCl (Thermo Fisher) for 30 min at 37°C before overnight incubation with primary antibody anti-BrdU (1:500; #ab6326, Abcam, Cambridge, UK) at 4°C. Secondary staining and mounting were performed as described above.

Image processing was performed using an LSM 780 confocal microscope (Zeiss, Jena, Germany) using 40x/1.4 Oil DIC objective. Four equally-spaced sections per mouse (rostral through the caudal extent of the DG) were used to count the total number of positive cells within the DG (subgranular zone and hilar subregions) of the hippocampus according to stereological principles. All counts were performed using FIJI in a blinded fashion, according to similar methods^4^.

#### Aggrecan and Homer1 puncta quantification

Quantification of the number of puncta by super-resolution microscopy proceeded similar to the method described previously^5^. Briefly, images were acquired by confocal microscopy imaging using an LSM 880 with AiryScan in super-resolution mode (Zeiss) set with a 63X/1.4 Oil DIC objective with 5X optical zoom. Using consistent laser strength, gain, and digital offset settings across all sections/experiments, Z stacks were acquired with steps less than the optical slices acquired. Using Zen software (Zeiss), a setting of 6 (“optimal”) was used during AiryScan processing. Aggrecan puncta were quantified using the Puncta Analyzer plugin^6^ in ImageJ, and thresholding was applied equally across images. For colocalization of co-stained Aggrecan and Homer1 puncta, images were analyzed with the Puncta Analyzer plugin with a minimum pixel specification of 4. Three images in the molecular layer of the DG were averaged per mouse for analysis.

#### Dendritic spine analysis by iontophoretic dye injections

*Tissue processing.* Mice were anesthetized using 15% chloral hydrate and transcardially perfused with 1% PFA in phosphate buffer (PB, pH 7.4) for 2 min, followed by 4% PFA with 0.125% glutaraldehyde in PB for 10 min at a rate of 5 ml/min. Brains were dissected and postfixed overnight at 4°C in the same fixative, and then transferred to PBS with 0.1% sodium azide at 4°C until sectioned. Brains were hemisected, and the right hemisphere was cut into 200 µm-thick sections using a vibratome (VT1000S, Leica).

*Intracellular dye injection and confocal imaging*. Coronal sections were incubated in 250 ng/ml DAPI for 5 min to enable identification of the DG. Sections were mounted on nitrocellulose membrane filters, immersed in ice-cold PB, and DG granule cells were iontophoretically injected with 5% Lucifer Yellow (Invitrogen, Waltham, MA, USA) in distilled water under a direct current of 3-8 nA until the dye filled the distal ends of the dendrites. Six to eleven neurons were injected per section and neurons selected for injection were spaced to avoid overlapping of dendrites. Sections were mounted with Fluoromount-G (#0100-01, Southern Biotech, Birmingham, AL, USA) between spacers placed on gelatin-coated glass slides (#22-214-320, Thermo Fisher).

To select dendritic segments for imaging on a Zeiss LSM 780 confocal microscope (Zeiss), whole cells at the suprapyramidal blade of the DG were first imaged using a 20x/0.8 M27 Plan-Apochromat objective, using an Ar/Kr laser at an excitation wavelength of 488 nm. Confocal stacks were acquired at 512 x 512-pixel resolution with a Z-step of 1 µm, a pinhole setting of 1 Airy Unit, and optimal settings for gain and offset. Basal dendritic segments at 50 µm from the soma were selected for high-resolution imaging according to the following criteria: not a primary dendrite, no overlap with other dendrites or branching that would obscure spines, not too deep in the section, and parallel or at acute angles to the coronal plane. Dendritic segments were imaged using a 100x/1.46 Oil DIC M27 Plan-Apochromat objective, and stacks were acquired at 512 x 512-pixel resolution with a Z-step of 0.1 µm, optical zoom of 3.3x, a pinhole setting of 1 Airy Unit, and optimal settings for gain and offset. The stacks were imaged with approximately 1 µm above and below the segment to fully include all spines. Three z-stacks were imaged from each neuron. Confocal stacks were deconvolved using an iterative blind deconvolution algorithm (AutoQuant X, vX3.0.1, MediaCybernetics, Rockville, MD, USA).

*Spine reconstruction and analysis.* Deconvolved stacks were analyzed using Neurolucida 360 (v2019.2.1; MBF Bioscience, Williston, VT, USA) for semi-automated reconstruction of dendrites and spines to determine spine density and morphology. Spines were classified as stubby, thin, mushroom, and filopodia, based on their morphologies, according to previous work^7^. 4-5 mice per genotype, 6 neurons per mouse, and 3 dendrites per neuron were analyzed, and violin plots were used to better visualize the distribution of individual data points.

#### Scanning electron microscopy (SEM)

Following ketamine/xylazine anesthesia and transcardial perfusion with PBS, 2-mm thick brain slices containing the hippocampus were decellularized, following previously described methods^8^. SEM methods were adapted from a previous study^9^. Decellularized brain slices were fixed in 4% PFA with 2% glutaraldehyde in 0.1M sodium cacodylate buffer (pH 7.4) for 24 hours at 4°C. Samples were then briefly rinsed in the same buffer before post-staining with 1% OsO4 for 1 hour. OsO4-treated samples were rinsed in water and gradually dehydrated in increasing concentrations of ethanol (50, 70, 90 100, and 100%, 10 min each). Samples were then stacked horizontally onto wire mesh dividers to keep them flattened and critical point-dried with liquid CO2. Dried samples were mounted onto Aluminum SEM stubs using conductive copper tape and sputter-coated before imaging with a Zeiss Supra 55 VP FESEM using InLens SE detection at 5 kV operating voltage (Zeiss Microscopy Inc). Fiber diameters were analyzed using Image J (Bethesda, USA).

#### Behavior

All behavioral analyses were performed during the 0700–1900 light cycle. For all behavioral experiments, age- (2-3-month-old) and sex-matched mice with corresponding littermate control animals were used.

*Novel location recognition.* Hippocampus-dependent memory was assessed using the novel location recognition as previously described^4^. On day 1, the training day, mice were habituated to the open-field arena for 6 min, an arena that contained wall-mounted visual cues. Mice were exposed to three consecutive trials of 6 min each, during which they explored two different objects in fixed positions. On day 2, the testing day, mice explored the same arena as on the training day but with one object displaced to a novel position. Time spent exploring each object was manually scored in a blinded fashion to assess the discrimination index for the novel location as the displaced object interaction time/total interaction time of both objects.

*Contextual fear-conditioning.* Fear-conditioning experiments were performed as previously described^4^ to assess hippocampus-dependent memory. Briefly, mice were trained to associate the cage context with an aversive stimulus (light foot-shock; Ugo Basile). On the first day, training parameters included two periods of 30 s consisting of a paired cue light and a tone of 1,000-Hz, followed by a light foot-shock (2 s, 0.5 mA) separated by a 180-s interval. Chambers were cleaned with 70% ethanol between experiments by the experimenter. Twenty-four hours later, mice were re-exposed to the same context for 3 min, and freezing levels (contextual) were measured. Two hours later, for the cued task, mice were placed in a novel context and exposed to the same tone and cue light from day 1 (training) after 120 s of exploration. Freezing levels were analyzed for pre- and post-cue phases. Freezing levels were measured using EthoVision XT system software (v14.0.1319, Noldus).

*Barnes maze.* Barnes maze testing was performed similarly to that previously described^4^. A large circular maze containing 40 holes was centered over a pedestal and elevated approximately 40 cm above the floor with a video recording device mounted directly above. The escape hole consisted of a PVC box similar in texture and color to the surface of the maze. Distinct visual cues were placed at four equally spaced points around the maze to serve as spatial navigation cues as the mice navigate the task. The task proceeded over 4 days, with four trials on each day for each mouse. With overhead illumination and a persistent 2 kHz tone, mice were given 90 s for each trial to identify the escape hole by jumping in or identifying the hole with extended/overhead pokes. If mice failed to find the escape hole within 90 s, they were gently guided by light tapping/directing towards the escape hole and scored with 90 s. The escape hole position was fixed within a day but changed for each successive day of testing. For each trial within a day, the starting location for the mouse was rotated relative to the escape hole position. Data collection and analyses were performed using EthoVision video-tracking system.

Search strategy classification was manually performed and based on methods adapted from previous work^10^, with data collected by EthoVision video-tracking system software. Search strategies were coded according to the following: (i) localized - minimal movement from the starting position; (ii) serial - sequential nose points within the outer ring area; (iii) random - unorganized searching; (iv) scanning - arc-like trajectory, without nose points; (v) focal - direct search within a maze quadrant; (vi) focal missense - direct search within a maze quadrant from previous trials; (vii) targeted - direct movement to hole with deviations in the trajectory direction; (viii) direct - no shifts in the trajectory direction with single nose point to target hole. For strategy analysis, each related strategy was categorized into non-hippocampus-dependent strategies comprising localized, serial, random, and scanning strategies, and hippocampus-dependent strategies that included focal, focal missense, targeted and direct strategies. Cognitive performance of each trial on day 3 was scored such that cognitive strategies received higher scores according to the following scale, accounting for high cognitive complexity, with a separation of 1 point between non-hippocampus-dependent and hippocampus-dependent strategies: localized = 0, serial = 1, random = 2, scanning = 3, focal = 5, focal missense = 5, targeted = 6, direct = 6. Focal and focal missense strategies are assigned a score of 5, as both are goal-directed strategies used without achieving the final target. Target and direct strategies were assigned score of 6 since both are goal-directed strategies with successful achievement of the final target.

*Open field*. The open field test was used to evaluate locomotion and exploratory behavior, and anxiety-like behavior in the center of the arena. The apparatus consisted of a brightly illuminated square arena of 50 x 50 cm enclosed by walls 45 cm high. Mice were placed individually in the center of the open field arena, and their movement was traced for 6 min. The resulting data was analyzed using EthoVision software, considering two previously defined areas: a central and an outer area. Due to the thigmotaxic exploratory activity of rodents, the ratio between the time spent in the center and in the periphery of the open field arena can reflect anxiety-like behaviors. Distance traveled, average velocity, and time spent in each of the zones were recorded and analyzed.

*Rotarod*. Balance and motor function was measured using the fixed and standard accelerated rotarod test (Med Associates). The protocol consisted of a first day of training at a constant speed (4 rpm) for a maximum of 2 min in three trials, with a 2-min interval between each trial. On day 2, animals were tested for each of 3 different fixed speeds over a range of 4 to 40 rpm for a maximum of 60 sec each in three trials, with a 2-min interval between each trial. For the acceleration test, animals were placed on the rod rotating at a constant speed (4 rpm), then the rod started to accelerate continuously from 4 to 40 rpm over 5 min, in three trials with a 5-min interval between each trial. The latency to fall off the rotarod was recorded for each trial, and data from three trials were averaged for each mouse.

*Wire grip test.* Motor coordination and strength was measured by the wire grip task. Each mouse was placed on a wire cage top, which was slowly inverted and suspended at approximately 30 cm above a mouse cage. The latency of time until the mouse fell off the wire was recorded in three trials. Mice that did not fall within the 60-s trial period were removed and assigned a maximal time of 60 s. Data from three trials were averaged for each mouse.

*Clasping.* Mice were suspended by their tail, and the extent of hindlimb clasping was observed for 15 s. Clasping scoring was based on a previously described method^11^. Briefly, if both hindlimbs were splayed outward away from the abdomen with splayed toes, a score of 0 was given. If one hindlimb was retracted or both hindlimbs were partially retracted toward the abdomen without touching it, and the toes were splayed, a score of 1 was assigned. If both hindlimbs were partially retracted toward the abdomen and were touching the abdomen without touching each other, a score of 2 was given. If both hindlimbs were fully clasped and touching the abdomen, a score of 3 was assigned.

*Grip strength.* For the grip strength test, forelimb, hindlimb, and 4-limbs grip strength was evaluated using a grip strength meter (Bioseb). Animals were held by the tail and allowed to place forelimbs on the bar, or they were restrained to place hindlimbs on the bar connected to the force sensor. Mice were slowly pulled away until they released the bar, and the maximal grip force measured during the pulling session was recorded. For the 4-limb measurement, using a mesh pull bar attachment, animals were handled by the tail so that all the paws latched onto the mesh pull the bar together. The mice were slowly and steadily pulled away from the apparatus exactly parallel from the countertop until paws were released from the mesh. A modified version of the forelimb grip strength test was performed as previously described^12^. Briefly, the meter was rotated vertically and the measurement procedure was identical to the conventional test, except for the direction in which the mouse’s tail was pulled. Each animal’s grip strength (in *g*) was recorded, and data from five trials were collected.

*Pole test.* Motor coordination and balance with striatal involvement was assessed with the vertical pole test, which consisted of a plastic pole (50 cm high, diameter 1 cm) placed vertically on a mouse cage. Mice were individually placed at the top of the pole facing upward by their front paws on the pole, and the time taken to turn around and climb down the pole was recorded. A maximum time of 120 s was given to complete the task. If the mouse fell from the top of the pole, a time of 120 s was recorded. Each mouse underwent three trials, and the pole was cleaned with 70% ethanol between animals. Data from three trials were averaged for each mouse.

#### qPCR

After rapid dissection of the hippocampus, tissue was preserved in RNAlater (Invitrogen) overnight at 4°C before storing at −80°C until use. Hippocampal RNA was extracted according to the manufacturer’s instructions using a RNeasy Mini Kit (Qiagen). Quality and concentration of total RNA were measured on the Nanodrop 8000 (Thermo Fisher). cDNA was subsequently synthesized with SuperScript III (Invitrogen), following the manufacturer’s recommendations. Samples were mixed with SYBR Green master mix (Thermo Fisher) and primers for *Mmp2* (FW: 5’-CAAGTTCCCCGGCGATGTC-3’; REV: 5’-TTCTGGTCAAGGTCACCTGTC-3’) before loading as technical replicates for qPCR on a QuantStudio 7 Flex Real-Time PCR System (Thermo Fisher). Cycle counts for mRNA quantification were normalized to *Rplp0* (FW: 5’- AGATTCGGGATATGCTGTTGGC-3’; REV: 5’-CCAGTTGGTAACAATGCCATGT-3’). The ΔΔCT method was used to provide gene expression values, and expression levels were normalized to *Rplp0*.

#### Western blotting

For Western blotting of MMP2 or TIMP2 within hippocampus, dissected hippocampi were manually homogenized with 60 strokes in RIPA lysis buffer (Thermo Fisher), with protease inhibitors (Roche), before centrifugation at 20,000g for 25 min at 4°C. Protein concentration of the supernatant was determined using the BCA kit (Thermo Fisher). Samples with equal amounts of protein were separated on 4-12% NuPAGE Bis-Tris precast denaturing gels (Invitrogen) and transferred onto nitrocellulose membranes (BioRad, Hercules, CA, USA) at 125 V for 90 min. Membranes were blocked with 3% milk-TBST (0.125% Tween) for 1 hour at room temperature and then probed with primary antibodies diluted in 3% milk-TBST solution overnight at 4°C: anti- TIMP2 (1:500; #D18B7, Cell Signaling), anti-MMP2 (1:1000; #ab86607, Abcam), and anti-actin (1:10000; #A5060, Sigma). Membranes were washed and probed with horseradish-peroxidase-conjugated anti-mouse (1:15,000; #1705047, BioRad) or anti-rabbit (1:15,000; #1705046, BioRad) for 1 hour at room temperature. Membranes were developed using ECL Clarity reagent (BioRad) and imaged using ChemiDoc MP Imaging System and Image Lab software (BioRad). Band intensities were quantified using ImageJ software as described previously^13^.

For TIMP2 detection in ISF samples, 4 consecutive baseline eISF samples from microdialysis were pooled and concentrated ∼6 fold using Amicon Ultra Centrifugal Filters (3 kDa Ultracel, 0.5 mL, Millipore). Concentrated samples were then processed for western blotting as described above. Estimation of exchangeable interstitial fluid (eISF) TIMP2 protein levels were determined by extrapolation from a standard curve of recombinant mouse TIMP2 (R&D/Biotechne) ranging from 0.31 ng to 20 ng run alongside ISF samples in SDS-PAGE/Western that had been concentrated ∼9.2-fold.

---

**Supplementary Figure 1.**
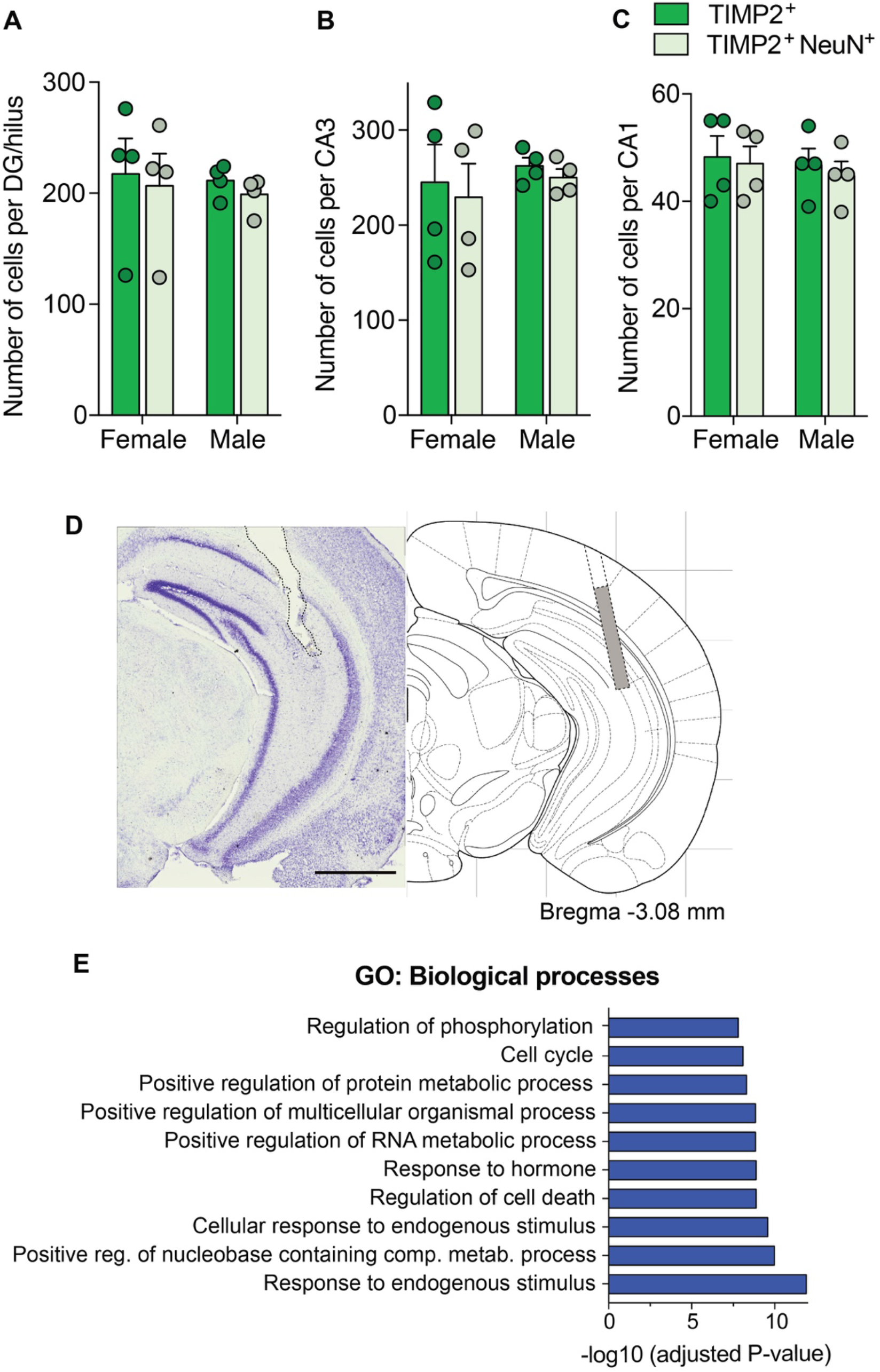
TIMP2 protein is found in hippocampal neurons in male and female mice, and its loss affects hippocampal transcriptome. (A) Quantification of the total number of TIMP2^+^ and TIMP2^+^ NeuN^+^ cells at the hilus/DG, (B) CA3 and (C) CA1 subregions of the hippocampus from WT males and females at 2 months of age (N = 4 mice per sex). **(D)** Cresyl violet-stained section depicting the microdialysis probe tract through brain surface extending into hippocampus, with microdialysis probe position seen at Bregma - 3.08mm based on Paxinos and Watson atlas. Scale bar, 500 μm. **(E)** Top 10 significant pathways for upregulated DEGs in hippocampi of TIMP2 KO mice relative to WT mice (N = 13-17 mice per group). Data are represented as mean ± SEM.

**Supplementary Figure S2.**
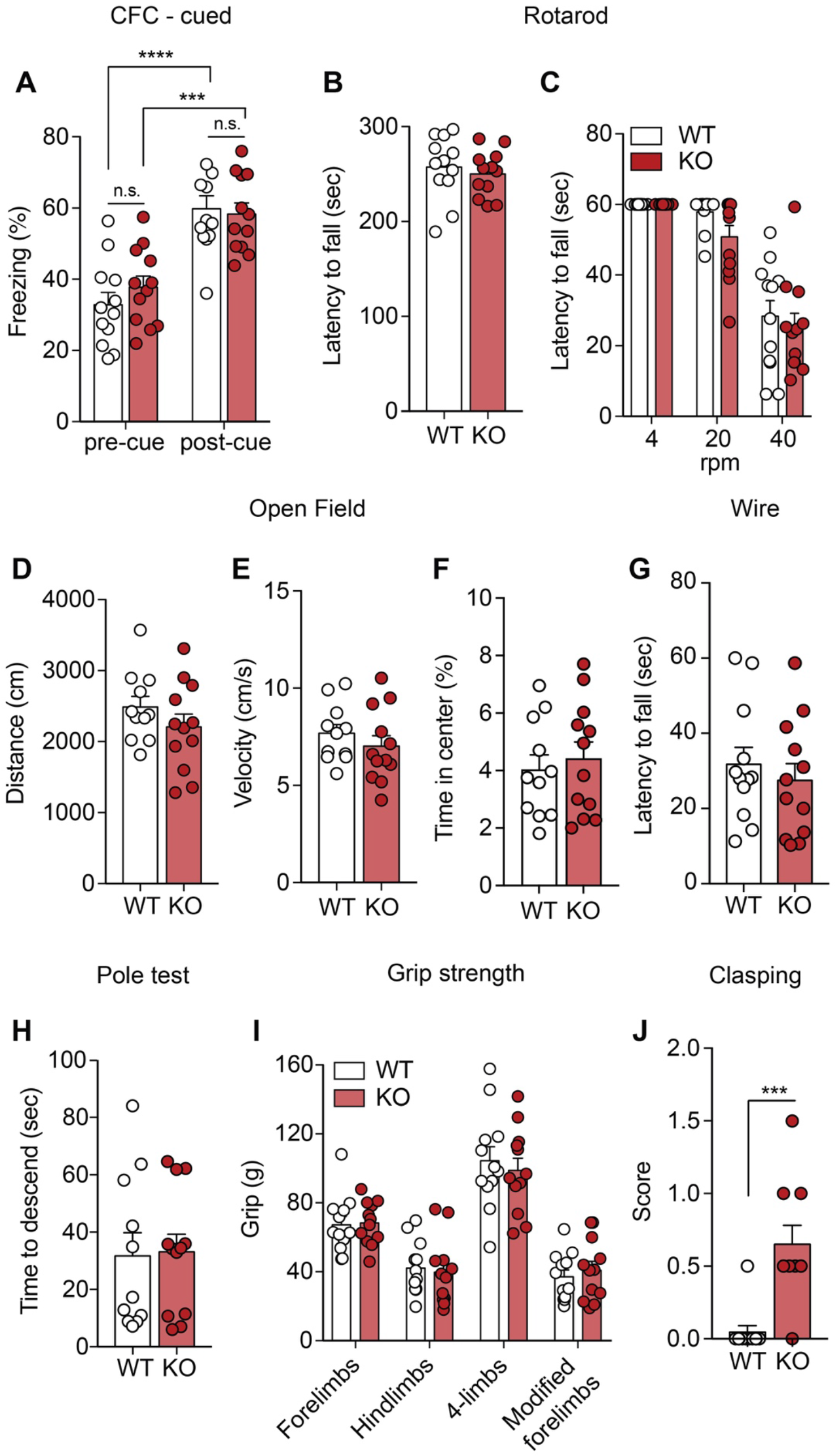
Loss of TIMP2 does not affect amygdala-associated behavior or overall motor function. (A) Percentage of freezing detected in the cued task of the fear-conditioning assay in WT and TIMP2 KO mice (2-3 months of age, N = 12 mice per group). **(B)** Latency of TIMP2 KO and WT mice to fall in the rotarod in the fixed and **(C)** acceleration protocol at 4, 20 and 40 rpm (N = 12 mice per group). **(D)** Total distance traveled by TIMP2 KO and WT mice in the open field, as well as **(E)** velocity, and **(F)** percentage of time spent in the center of the arena (N = 11-12 mice per group). **(G)** Latency of TIMP2 KO and WT mice to fall in the wire test (N = 12 mice per group). **(H)** Time for TIMP2 KO and WT mice to descend the pole in the pole test (N = 11-12 mice per group). **(J)** Grip strength of the fore-, hind- and four limbs in TIMP2 KO and WT mice (N = 12 mice per group). **(J)** Hindlimb extension by clasping score in TIMP2 KO and WT mice (N = 10-11 mice per group). Data are represented as mean ± SEM. Student’s *t*-test for two-group comparisons. n.s., not significant.

**Supplementary Figure 3.**
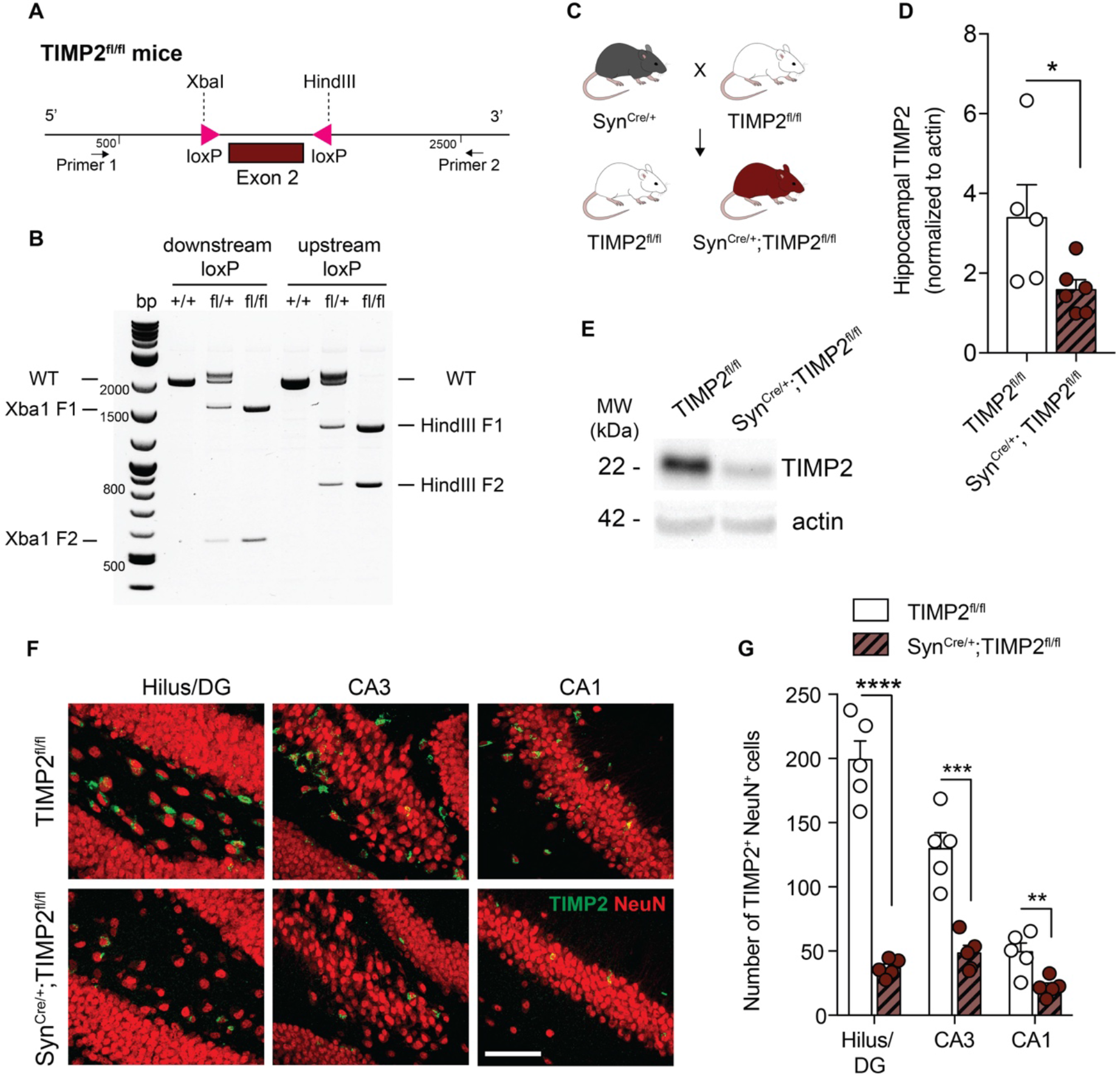
Creation of a conditional model to target neuronal pools of TIMP2. **(A)** Schematic illustration of the targeting strategy of to insert loxP sites flanking exon 2 to generate a model for conditional deletion of *TIMP2*. **(B)** Representative genotyping results revealing the PCR products for the mutant LoxP and wild-type alleles. Wild-type (+/+), heterozygous (fl/+), and homozygous (fl/fl) mice were identified according to this strategy. **(C)** Schematic diagram of cross-breeding strategy to establish neuron-specific TIMP2 deletion: male TIMP2^fl/fl^ mice were mated with Syn^Cre/+^ females to obtain Syn^Cre/+^; TIMP2^fl/fl^ and their respective TIMP2^fl/fl^ littermate controls. **(D-E)** Representative TIMP2 immunoblot and corresponding quantification of TIMP2 protein levels from hippocampal lysate of TIMP2^fl/fl^ and Syn^Cre/+^; TIMP2^fl/fl^ (2-3 months of age, N = 5-6 mice per group). **(F)** High-magnification view of hilus/DG, CA3, and CA1 sub-regions of TIMP2+ cells co-expressing NeuN in TIMP2^fl/fl^ and Syn^Cre/+^; TIMP2^fl/fl^ mice (2- 3 months of age, N = 5 mice per group, scale bar, 20 μm) with corresponding **(G)** quantification of the total number of TIMP2^+^ cells with NeuN^+^ nuclei across hippocampal subregions. Data are represented as mean ± SEM. Student’s t-test for two-group comparisons. \**P*<0.05, \*\**P*<0.01, \*\*\**P*<0.001, \*\*\*\**P*<0.0001. Data points represent individual mice.

**Supplementary Figure 4.**
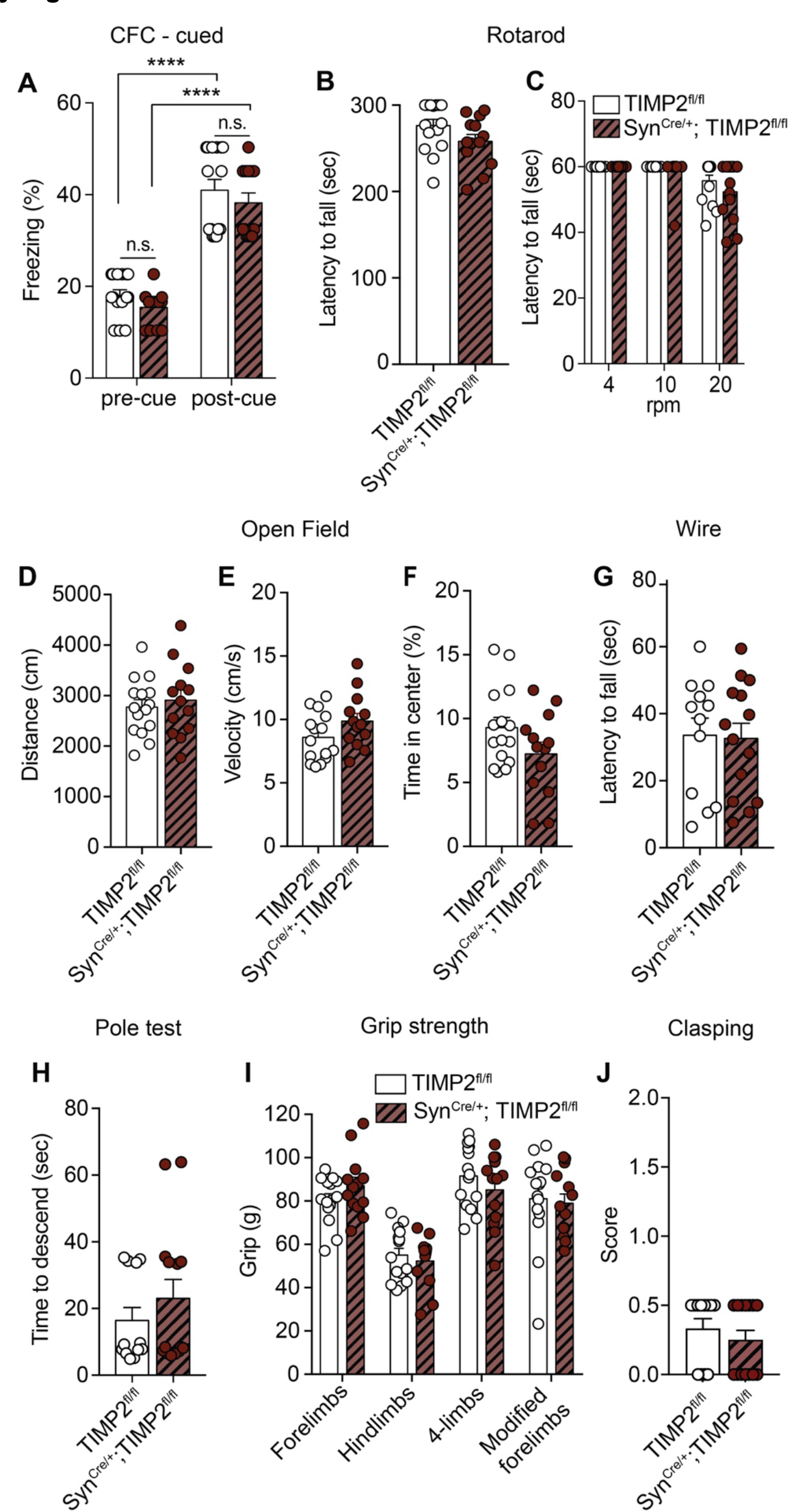
Neuronal TIMP2 deletion does not impair amygdala-associated behavior or overall motor function. **(A)** Percentage of freezing detected in the cued task of the fear-conditioning assay in TIMP2^fl/fl^ and Syn^Cre/+^; TIMP2^fl/fl^ mice (2-3 months of age, N = 13-15 mice per group). **(B)** Latency of TIMP2^fl/fl^ and Syn^Cre/+^; TIMP2^fl/fl^ mice to fall in the rotarod in the fixed and **(C)** acceleration protocol at 4, 10 and 20 rpm (N = 13-15 mice per group). **(D)** Total distance traveled by TIMP2^fl/fl^ and Syn^Cre/+^; TIMP2^fl/fl^ mice in the open field, as well as **(E)** velocity, and **(F)** percentage of time spent in the center of the arena (N = 13-15 mice per group). **(G)** Latency of TIMP2^fl/fl^ and Syn^Cre/+^; TIMP2^fl/fl^ mice to fall in the wire test (N = 12-14 mice per group). **(H)** Time for TIMP2^fl/fl^ and Syn^Cre/+^; TIMP2^fl/fl^ mice to descend the pole in the pole test (N = 12-14 mice per group). **(I)** Grip strength of the fore-, hind- and four limbs in TIMP2^fl/fl^ and Syn^Cre/+^; TIMP2^fl/fl^ mice (N = 13-15 mice per group). **(K)** Hindlimb extension by clasping score in TIMP2^fl/fl^ and Syn^Cre/+^; TIMP2^fl/fl^ mice (N = 12-14 mice per group). Data are represented as mean ± SEM. Student’s *t*-test for two-group comparisons. n.s., not significant.

## Notes

### Summary of Updates

Here we provide new data in Figure 5, updates in Figure 2, and additional methods details.

## References

1. Villeda SA, Luo J, Mosher KI, Zou B, Britschgi M, Bieri G et al. The ageing systemic milieu negatively regulates neurogenesis and cognitive function. Nature 2011; 477(7362): 90–94.

2. Villeda SA, Plambeck KE, Middeldorp J, Castellano JM, Mosher KI, Luo J et al. Young blood reverses age-related impairments in cognitive function and synaptic plasticity in mice. Nat Med 2014; 20(6): 659–663.

3. Katsimpardi L, Litterman NK, Schein PA, Miller CM, Loffredo FS, Wojtkiewicz GR et al. Vascular and neurogenic rejuvenation of the aging mouse brain by young systemic factors. Science 2014; 344(6184): 630–634.

4. Castellano JM, Mosher KI, Abbey RJ, McBride AA, James ML, Berdnik D et al. Human umbilical cord plasma proteins revitalize hippocampal function in aged mice. Nature 2017; 544(7651): 488–492.

5. Brew K, Nagase H. The tissue inhibitors of metalloproteinases (TIMPs): an ancient family with structural and functional diversity. Biochim Biophys Acta 2010; 1803(1): 55–71.

6. Caterina JJ, Yamada S, Caterina NC, Longenecker G, Holmback K, Shi J et al. Inactivating mutation of the mouse tissue inhibitor of metalloproteinases-2(Timp-2) gene alters proMMP-2 activation. J Biol Chem 2000; 275(34): 26416–26422.

7. Wang Z, Juttermann R, Soloway PD. TIMP-2 is required for efficient activation of proMMP- 2 in vivo. J Biol Chem 2000; 275(34): 26411–26415.

8. Kandalam V, Basu R, Moore L, Fan D, Wang X, Jaworski DM et al. Lack of tissue inhibitor of metalloproteinases 2 leads to exacerbated left ventricular dysfunction and adverse extracellular matrix remodeling in response to biomechanical stress. Circulation 2011; 124(19): 2094–2105.

9. Kaczorowska A, Miekus N, Stefanowicz J, Adamkiewicz-Drozynska E. Selected Matrix Metalloproteinases (MMP-2, MMP-7) and Their Inhibitor (TIMP-2) in Adult and Pediatric Cancer. Diagnostics (Basel) 2020; 10(8).

10. Guo T, Hao H, Zhou L, Zhou F, Yu D. Association of SNPs in the TIMP-2 gene and large artery atherosclerotic stroke in southern Chinese Han population. Oncotarget 2018; 9(4): 4698–4706.

11. Fager N, Jaworski DM. Differential spatial distribution and temporal regulation of tissue inhibitor of metalloproteinase mRNA expression during rat central nervous system development. Mech Dev 2000; 98(1-2): 105–109.

12. Perez-Martinez L, Jaworski DM. Tissue inhibitor of metalloproteinase-2 promotes neuronal differentiation by acting as an anti-mitogenic signal. J Neurosci 2005; 25(20): 4917–4929.

13. Jaworski DM, Soloway P, Caterina J, Falls WA. Tissue inhibitor of metalloproteinase- 2(TIMP-2)-deficient mice display motor deficits. J Neurobiol 2006; 66(1): 82–94.

14. Nicaise AM, Johnson KM, Willis CM, Guzzo RM, Crocker SJ. TIMP-1 Promotes Oligodendrocyte Differentiation Through Receptor-Mediated Signaling. Mol Neurobiol 2019; 56(5): 3380–3392.

15. Dewing JM, Carare RO, Lotery AJ, Ratnayaka JA. The Diverse Roles of TIMP-3: Insights into Degenerative Diseases of the Senescent Retina and Brain. Cells 2019; 9(1).

16. Solga R, Behrens J, Ziemann A, Riou A, Berwanger C, Becker L et al. CRN2 binds to TIMP4 and MMP14 and promotes perivascular invasion of glioblastoma cells. Eur J Cell Biol 2019; 98(5-8): 151046.

17. Ulrich JD, Burchett JM, Restivo JL, Schuler DR, Verghese PB, Mahan TE et al. In vivo measurement of apolipoprotein E from the brain interstitial fluid using microdialysis. Mol Neurodegener 2013; 8: 13.

18. Nguyen PT, Dorman LC, Pan S, Vainchtein ID, Han RT, Nakao-Inoue H et al. Microglial Remodeling of the Extracellular Matrix Promotes Synapse Plasticity. Cell 2020; 182(2): 388–403 e315.

19. Ippolito DM, Eroglu C. Quantifying synapses: an immunocytochemistry-based assay to quantify synapse number. J Vis Exp 2010; (45).

20. Jacot-Descombes S, Keshav NU, Dickstein DL, Wicinski B, Janssen WGM, Hiester LL et al. Altered synaptic ultrastructure in the prefrontal cortex of Shank3-deficient rats. Mol Autism 2020; 11(1): 89.

21. Dickstein DL, Dickstein DR, Janssen WGM, Hof PR, Glaser JR, Rodriguez A et al. Automatic Dendritic Spine Quantification from Confocal Data with Neurolucida 360. Curr Protoc Neurosci 2016; 77: 1 27 21–21 27 21.

22. Mateus-Pinheiro A, Alves ND, Patricio P, Machado-Santos AR, Loureiro-Campos E, Silva JM et al. AP2gamma controls adult hippocampal neurogenesis and modulates cognitive, but not anxiety or depressive-like behavior. Mol Psychiatry 2017; 22(12): 1725–1734.

23. Yang G, Pan F, Gan WB. Stably maintained dendritic spines are associated with lifelong memories. Nature 2009; 462(7275): 920–924.

24. Price KA, Varghese M, Sowa A, Yuk F, Brautigam H, Ehrlich ME et al. Altered synaptic structure in the hippocampus in a mouse model of Alzheimer’s disease with soluble amyloid-beta oligomers and no plaque pathology. Mol Neurodegener 2014; 9: 41.

25. Rodriguez A, Ehlenberger DB, Dickstein DL, Hof PR, Wearne SL. Automated three- dimensional detection and shape classification of dendritic spines from fluorescence microscopy images. PLoS One 2008; 3(4): e1997.

26. Bourne J, Harris KM. Do thin spines learn to be mushroom spines that remember? Curr Opin Neurobiol 2007; 17(3): 381–386.

27. Zuo Y, Lin A, Chang P, Gan WB. Development of long-term dendritic spine stability in diverse regions of cerebral cortex. Neuron 2005; 46(2): 181–189.

28. Anacker C, Hen R. Adult hippocampal neurogenesis and cognitive flexibility - linking memory and mood. Nat Rev Neurosci 2017; 18(6): 335–346.

29. Jedlicka P, Vlachos A, Schwarzacher SW, Deller T. A role for the spine apparatus in LTP and spatial learning. Behav Brain Res 2008; 192(1): 12–19.

30. Moser MB, Trommald M, Andersen P. An increase in dendritic spine density on hippocampal CA1 pyramidal cells following spatial learning in adult rats suggests the formation of new synapses. Proc Natl Acad Sci U S A 1994; 91(26): 12673–12675.

31. Illouz T, Madar R, Louzoun Y, Griffioen KJ, Okun E. Unraveling cognitive traits using the Morris water maze unbiased strategy classification (MUST-C) algorithm. Brain Behav Immun 2016; 52: 132–144.

32. Magnowska M, Gorkiewicz T, Suska A, Wawrzyniak M, Rutkowska-Wlodarczyk I, Kaczmarek L et al. Transient ECM protease activity promotes synaptic plasticity. Sci Rep 2016; 6: 27757.

33. Yamada J, Nadanaka S, Kitagawa H, Takeuchi K, Jinno S. Increased Synthesis of Chondroitin Sulfate Proteoglycan Promotes Adult Hippocampal Neurogenesis in Response to Enriched Environment. J Neurosci 2018; 38(39): 8496–8513.

34. Vegh MJ, Rausell A, Loos M, Heldring CM, Jurkowski W, van Nierop P et al. Hippocampal extracellular matrix levels and stochasticity in synaptic protein expression increase with age and are associated with age-dependent cognitive decline. Mol Cell Proteomics 2014; 13(11): 2975–2985.

35. Miyata S, Nadanaka S, Igarashi M, Kitagawa H. Structural Variation of Chondroitin Sulfate Chains Contributes to the Molecular Heterogeneity of Perineuronal Nets. Front Integr Neurosci 2018; 12: 3.

36. Zimmermann DR, Dours-Zimmermann MT. Extracellular matrix of the central nervous system: from neglect to challenge. Histochem Cell Biol 2008; 130(4): 635–653.

37. Rowlands D, Lensjo KK, Dinh T, Yang S, Andrews MR, Hafting T et al. Aggrecan Directs Extracellular Matrix-Mediated Neuronal Plasticity. J Neurosci 2018; 38(47): 10102–10113.

38. Fawcett JW, Oohashi T, Pizzorusso T. The roles of perineuronal nets and the perinodal extracellular matrix in neuronal function. Nat Rev Neurosci 2019; 20(8): 451–465.

39. Gogolla N, Caroni P, Luthi A, Herry C. Perineuronal nets protect fear memories from erasure. Science 2009; 325(5945): 1258–1261.

40. Romberg C, Yang S, Melani R, Andrews MR, Horner AE, Spillantini MG et al. Depletion of perineuronal nets enhances recognition memory and long-term depression in the perirhinal cortex. J Neurosci 2013; 33(16): 7057–7065.

41. Cope EC, Gould E. Adult Neurogenesis, Glia, and the Extracellular Matrix. Cell Stem Cell 2019; 24(5): 690–705.

42. Dansie LE, Ethell IM. Casting a net on dendritic spines: the extracellular matrix and its receptors. Dev Neurobiol 2011; 71(11): 956–981.

43. Singhal N, Martin PT. Role of extracellular matrix proteins and their receptors in the development of the vertebrate neuromuscular junction. Dev Neurobiol 2011; 71(11): 982–1005.

44. Jaworski DM, Perez-Martinez L. Tissue inhibitor of metalloproteinase-2 (TIMP-2) expression is regulated by multiple neural differentiation signals. J Neurochem 2006; 98(1): 234–247.

45. Wang S, Brunne B, Zhao S, Chai X, Li J, Lau J et al. Trajectory Analysis Unveils Reelin’s Role in the Directed Migration of Granule Cells in the Dentate Gyrus. J Neurosci 2018; 38(1): 137–148.

46. Levy AD, Omar MH, Koleske AJ. Extracellular matrix control of dendritic spine and synapse structure and plasticity in adulthood. Front Neuroanat 2014; 8: 116.

47. Choi YS, Vincent LG, Lee AR, Kretchmer KC, Chirasatitsin S, Dobke MK et al. The alignment and fusion assembly of adipose-derived stem cells on mechanically patterned matrices. Biomaterials 2012; 33(29): 6943–6951.

48. Vincent LG, Choi YS, Alonso-Latorre B, del Alamo JC, Engler AJ. Mesenchymal stem cell durotaxis depends on substrate stiffness gradient strength. Biotechnol J 2013; 8(4): 472–484.

49. Saha K, Keung AJ, Irwin EF, Li Y, Little L, Schaffer DV et al. Substrate modulus directs neural stem cell behavior. Biophys J 2008; 95(9): 4426–4438.

50. Leipzig ND, Shoichet MS. The effect of substrate stiffness on adult neural stem cell behavior. Biomaterials 2009; 30(36): 6867–6878.

51. Rammensee S, Kang MS, Georgiou K, Kumar S, Schaffer DV. Dynamics of Mechanosensitive Neural Stem Cell Differentiation. Stem Cells 2017; 35(2): 497–506.

52. Barnes JM, Przybyla L, Weaver VM. Tissue mechanics regulate brain development, homeostasis and disease. J Cell Sci 2017; 130(1): 71–82.

53. Hall CM, Moeendarbary E, Sheridan GK. Mechanobiology of the brain in ageing and Alzheimer’s disease. Eur J Neurosci 2021; 53(12): 3851–3878.

54. Javier-Torrent M, Zimmer-Bensch G, Nguyen L. Mechanical Forces Orchestrate Brain Development. Trends Neurosci 2021; 44(2): 110–121.

55. Keren-Shaul H, Spinrad A, Weiner A, Matcovitch-Natan O, Dvir-Szternfeld R, Ulland TK et al. A Unique Microglia Type Associated with Restricting Development of Alzheimer’s Disease. Cell 2017; 169(7): 1276–1290 e1217.

56. Birch HL. Extracellular Matrix and Ageing. Subcell Biochem 2018; 90: 169–190.

57. Sun Y, Xu S, Jiang M, Liu X, Yang L, Bai Z et al. Role of the Extracellular Matrix in Alzheimer’s Disease. Front Aging Neurosci 2021; 13: 707466.

58. Bettio LEB, Rajendran L, Gil-Mohapel J. The effects of aging in the hippocampus and cognitive decline. Neurosci Biobehav Rev 2017; 79: 66–86.

59. Vegh MJ, Heldring CM, Kamphuis W, Hijazi S, Timmerman AJ, Li KW et al. Reducing hippocampal extracellular matrix reverses early memory deficits in a mouse model of Alzheimer’s disease. Acta Neuropathol Commun 2014; 2: 76.

60. Moreno-Jimenez EP, Flor-Garcia M, Terreros-Roncal J, Rabano A, Cafini F, Pallas- Bazarra N et al. Adult hippocampal neurogenesis is abundant in neurologically healthy subjects and drops sharply in patients with Alzheimer’s disease. Nat Med 2019; 25(4): 554–560.

## References

1. Gu B, Posfai E, Rossant J. Efficient generation of targeted large insertions by microinjection into two-cell-stage mouse embryos. Nat Biotechnol 2018; 36(7): 632–637.

2. Zhu Y, Romero MI, Ghosh P, Ye Z, Charnay P, Rushing EJ et al. Ablation of NF1 function in neurons induces abnormal development of cerebral cortex and reactive gliosis in the brain. Genes Dev 2001; 15(7): 859–876.

3. Ulrich JD, Burchett JM, Restivo JL, Schuler DR, Verghese PB, Mahan TE et al. In vivo measurement of apolipoprotein E from the brain interstitial fluid using microdialysis. Mol Neurodegener 2013; 8: 13.

5. Nguyen PT, Dorman LC, Pan S, Vainchtein ID, Han RT, Nakao-Inoue H et al. Microglial Remodeling of the Extracellular Matrix Promotes Synapse Plasticity. Cell 2020; 182(2): 388–403 e315.

6. Ippolito DM, Eroglu C. Quantifying synapses: an immunocytochemistry-based assay to quantify synapse number. J Vis Exp 2010; (45).

7. Jacot-Descombes S, Keshav NU, Dickstein DL, Wicinski B, Janssen WGM, Hiester LL et al. Altered synaptic ultrastructure in the prefrontal cortex of Shank3-deficient rats. Mol Autism 2020; 11(1): 89.

8. De Waele J, Reekmans K, Daans J, Goossens H, Berneman Z, Ponsaerts P. 3D culture of murine neural stem cells on decellularized mouse brain sections. Biomaterials 2015; 41: 122–131.

9. Tajerian M, Hung V, Nguyen H, Lee G, Joubert LM, Malkovskiy AV et al. The hippocampal extracellular matrix regulates pain and memory after injury. Mol Psychiatry 2018; 23(12): 2302–2313.

10. Mateus-Pinheiro A, Alves ND, Patricio P, Machado-Santos AR, Loureiro-Campos E, Silva JM et al. AP2gamma controls adult hippocampal neurogenesis and modulates cognitive, but not anxiety or depressive-like behavior. Mol Psychiatry 2017; 22(12): 1725–1734.

11. Zhu JW, Li YF, Wang ZT, Jia WQ, Xu RX. Toll-Like Receptor 4 Deficiency Impairs Motor Coordination. Front Neurosci 2016; 10: 33.

12. Takeshita H, Yamamoto K, Nozato S, Inagaki T, Tsuchimochi H, Shirai M et al. Modified forelimb grip strength test detects aging-associated physiological decline in skeletal muscle function in male mice. Sci Rep 2017; 7: 42323.

13. Castellano JM, Kim J, Stewart FR, Jiang H, DeMattos RB, Patterson BW et al. Human apoE isoforms differentially regulate brain amyloid-beta peptide clearance. Sci Transl Med 2011; 3(89): 89ra57.

